# Istaroxime metabolite PST3093 selectively stimulates SERCA2a and reverses disease-induced changes in cardiac function

**DOI:** 10.1101/2021.08.17.455204

**Authors:** Martina Arici, Mara Ferrandi, Paolo Barassi, Shih-Che Hsu, Eleonora Torre, Andrea Luraghi, Carlotta Ronchi, Gwo-Jyh Chang, Francesco Peri, Patrizia Ferrari, Giuseppe Bianchi, Marcella Rocchetti, Antonio Zaza

**Author notes:** Correspondence: AZ; MR. MA and MF contributed equally as first authors to the article. MR and AZ contributed equally as senior authors to the article.

## Abstract

**Background:** Heart failure (HF) therapeutic toolkit would strongly benefit from the availability of ino-lusitropic agents with a favorable pharmacodynamics and safety profile. Istaroxime is a promising agent, which combines Na^+^/K^+^ pump inhibition with SERCA2a stimulation; however, it has a very short half-life and extensive metabolism to a molecule, named PST3093. The present work aims to investigate whether PST3093, still retains the pharmacodynamic and pharmacokinetic properties of its parent compound.

**Methods:** We studied PST3093 for its effects on SERCA2a and Na^+^/K^+^ ATPase activities, Ca^2+^ dynamics in isolated myocytes and hemodynamic effects in an *in-vivo* rat model of diabetic (streptozotocin (STZ)-induced) cardiomyopathy.

**Results:** Istaroxime infusion in HF patients led to accumulation of PST3093 in the plasma; clearance was substantially slower for PST3093 than for istaroxime. In cardiac rat preparations PST3093 did not inhibit the Na^+^/K^+^ ATPase activity, but retained SERCA2a stimulatory activity. In *in-vivo* echocardiographic assessment, PST3093 improved overall cardiac performance and reversed most STZ-induced abnormalities. PST3093 i.v. toxicity was considerably lower than that of istaroxime and it failed to significantly interact with 50 off-targets.

**Conclusions:** Overall, PST3093 is a “selective” SERCA2a activator, the prototype of a novel pharmacodynamic category with a potential in the ino-lusitropic approach to HF with prevailing diastolic dysfunction. Its pharmacodynamics are peculiar and its pharmacokinetics are suitable to prolong the cardiac beneficial effect of istaroxime infusion.

**Graphical abstract:** 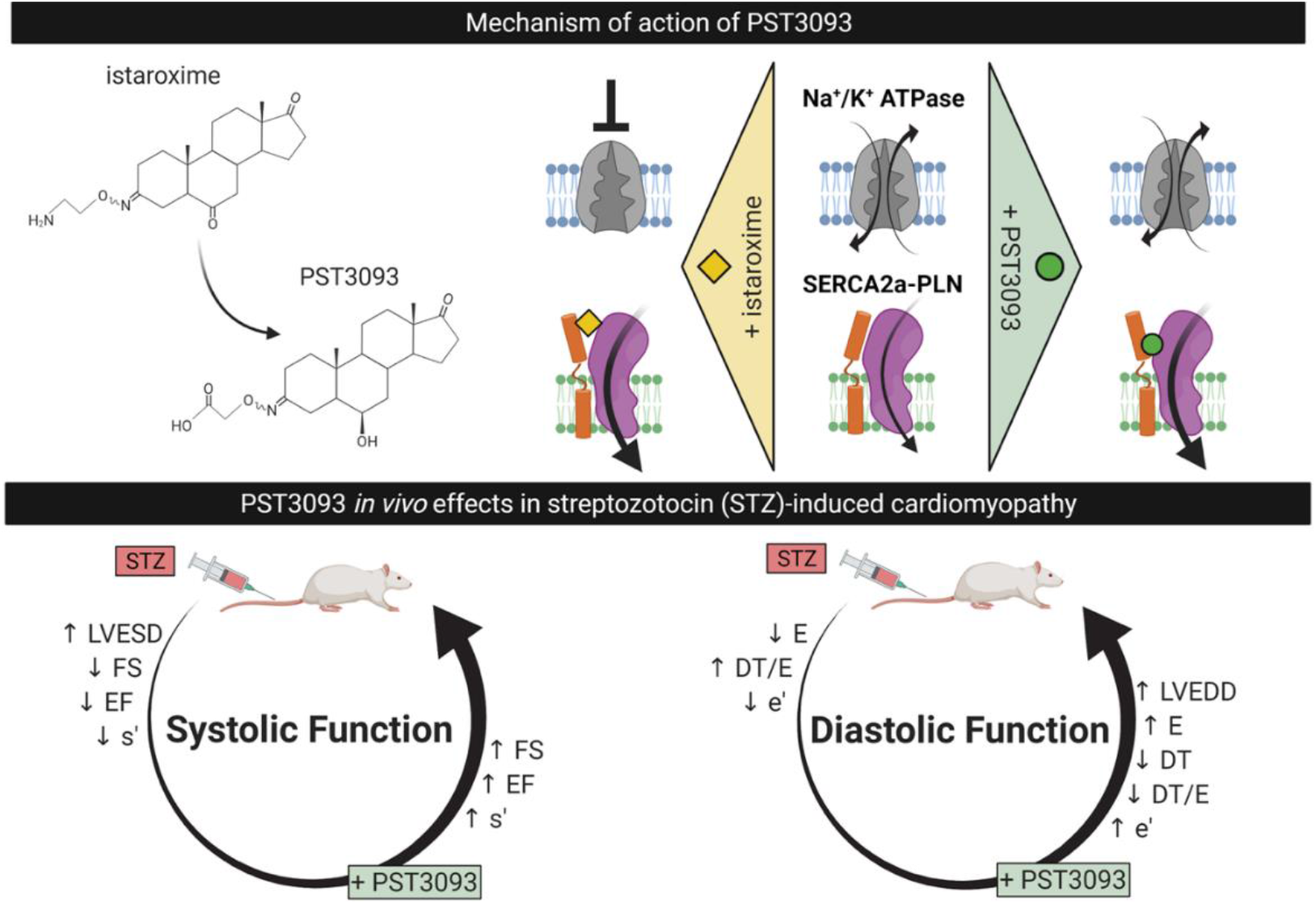

## Introduction

Heart failure (HF) is characterized by abnormal Ca^2+^ distribution among subcellular compartments, which contributes to impaired contractility and relaxation (Bers and Despa, 2006), facilitates arrhythmias (Zaza and Rocchetti, 2015) and, in the long run, contributes to myocardial remodeling (Nakayama et al., 2007). Evidence of a deficient SERCA2a activity in HF dates to the 70’s (Suko et al., 1970; Sulakhe and Dhalia, 1971). Since then, many studies confirmed this finding (Arai et al., 1993; Gwathmey et al., 1987; Kranias and Hajjar, 2012) showing that the impaired SERCA2a activity can often result from an over-inhibition by phospholamban (PLN) (Del Monte et al., 2002; Haghighi et al., 2001). Loss of SERCA2a function accounts for abnormal distribution of intracellular Ca^2+^ with numerous detrimental consequences. Interventions currently available to modulate myocyte Ca^2+^ handling (e.g. amines, PDE inhibitors etc.) stimulate SERCA2a, but they do so in the context of a multi-target action, thus resulting in untoward effects. Selective SERCA2a enhancement would afford inotropic and lusitropic effects without the drawbacks of the multi-target action (Zaza and Rocchetti, 2015). Accordingly, the use of SERCA2a stimulation in HF therapy is receiving considerable attention and many attempts to selectively stimulate SERCA2a activity with gene therapy or “small molecule” agents have been reported (Kaneko et al., 2017; Kho et al., 2011; Schaaf et al., 2020). Nonetheless, for reasons other than refutation of the principle, none of these attempts has been successfully translated into the clinic. The only exception is istaroxime, a small-molecule drug, identified as SERCA2a enhancer by our group (Rocchetti et al., 2005) and currently under clinical development for the treatment of acute HF (Carubelli et al., 2020; Shah et al., 2009). Istaroxime has a double mechanism of action: it inhibits the Na^+^/K^+^ pump (Micheletti et al., 2002) and activates SERCA2a (Rocchetti et al., 2005). Thus, istaroxime increases overall cell Ca^2+^ content while promoting rapid Ca^2+^ sequestration into the sarcoplasmic reticulum (SR). Notably, at variance with Na^+^/K^+^ pump blockade alone, this neither facilitates spontaneous Ca^2+^ release from the SR (Alemanni et al., 2011), nor increases myocardial oxygen demand (Sabbah et al., 2007). Thus, istaroxime may improve systolic and diastolic performance (Shah et al., 2009) without promoting arrhythmia or ischemia (Carubelli et al., 2020; Gheorghiade et al., 2008). However, istaroxime has a plasma half-life of less than 1 hour, because of extensive hepatic metabolism to a molecule, named PST3093 (Carubelli et al., 2020; Gheorghiade et al., 2008); this restricts istaroxime usage to acute intravenous therapy.

The present work aims to investigate whether PST3093 may, on its own, be endowed with pharmacological activity. To this end, PST3093 has been synthesized and compared to istaroxime and digoxin (as reference compounds) in experimental setups at different levels of biological organization and in the context of disease-induced dysfunction.

The data here reported indicate that PST3093 shows a longer half-life in human circulation compared to parent drug, it stimulates SERCA2a activity but, at variance with istaroxime it does not inhibit the Na^+^/K^+^ ATPase. This pharmacodynamic profile translates to positive inotropy and lusitropy in an *in-vivo* disease model characterized by SERCA2a downregulation. Therefore, PST3093 qualifies as a “pure” SERCA2a activator, able to improve cardiac mechanical performance in-vivo.

## Methods

### Further detail on methods is given in the Online Supplement

#### Pharmacokinetics

Pharmacokinetics (PKs) of istaroxime and its metabolite PST3093 was assessed in 30 heart failure (HF) patients infused for 6 hours with istaroxime at 1.0 μg/kg/min (secondary analysis of the HORIZON-HF study, NCT00616161) (Gheorghiade et al., 2008). Blood samples were taken before, during, and up to 18 hours after starting the infusion. The lowest concentration resolved by the technique was 2.6 ng mL^-1^ for istaroxime and 2.9 ng mL^-1^ for PST3093, lower values were considered as zero. PKs parameters were estimated using the dedicated software Kinetica (version 4.4, Thermo Electron Corp., Waltham, Massachusetts). Samples were excluded from the analysis if contamination was suspected, or ≥ 2 consecutive samples were missing. The following PKs parameters were estimated: maximum observed concentration (C_max_), the time of maximum observed concentration (T_max_), the elimination half life time (T_0.5_) and the area under the concentration curve from the start of istaroxime administration to the time of final sampling (AUC_last_).

#### Animal model

The animal study protocols were approved by the Institutional Review Board of Milano Bicocca (29C09.26 and 29C09.N.YRR protocol codes approved on January 2021 and June 2018 respectively) and Chang Gung (CGU107-068 protocol code approved on June 2018) Universities according to the Directive 2010/63/EU of the European Parliament on the protection of animals used for scientific purposes.

Streptozotocin (STZ)-induced diabetes was selected as a pathological model because of its association with reduced SERCA2a function (Torre et al., 2021) and relevance to diastolic dysfunction (Valero-Muñoz et al., 2017), for which a lusitropic action may be more relevant. Diabetes was induced in Sprague Dawley male rats (150-175 g) by a single STZ (50 mg.kg^-1^) i.v. injection (STZ group). Control (healthy group) rats received STZ vehicle (citrate buffer). Fasting glycaemia was measured after 1 week and rats with values >290 mg.dL^-1^ were considered diabetic (Torre et al., 2021). Rats were euthanized by cervical dislocation under anaesthesia with ketamine-xylazine (130-7.5 mg kg^-1^ i.p) 9 weeks after STZ injection.

#### Biochemical measurements

Total ATPase activity was assessed by measuring the rate of ^32^P-ATP release (μmol.min^-1^) at 37°C.

### Na^+^/K^+^ ATPase activity assay

The inhibitory effect of compounds was tested, at multiple concentrations, on suspensions of the enzyme α1 isoform from dog kidney by measuring ^32^P-ATP hydrolysis as previously described (Ferrandi et al., 1996). Na^+^/K^+^ ATPase activity was identified as the ouabain (1 mM)-sensitive component of total one; compound efficacy was expressed as the concentration exerting 50% inhibition (IC_50_).

### SERCA ATPase activity assay

Measurements were performed in whole tissue homogenates (rat) or in cardiac SR enriched microsomes (guinea-pig), including SERCA2a and PLN (Torre et al., 2021). To test for PLN involvement in the effect of compounds, SERCA1 activity was also measured in PLN-free microsomes (from guinea-pig skeletal muscle) before and after reconstitution with the PLN_1-32_ inhibitory fragment at a ratio PLN:SERCA of 300:1. The SERCA component, identified as the cyclopiazonic acid (CPA, 10 µM)-sensitive one, was measured at multiple Ca^2+^ concentrations (100-2000 nM) as ^32^P-ATP hydrolysis (Micheletti et al., 2007) and Ca^2+^ dose-response curves were fitted to estimate SERCA maximal hydrolytic velocity (V_max,_ μmol·min^-1.^mg^-1^ protein) and Ca^2+^ dissociation constant (K_d_Ca, nM). Either an increase of V_max_, or a decrease of K_d_Ca (increased Ca^2+^ affinity) stand for enhancement of SERCA function.

#### PST3093 interaction with targets other than SERCA

To predict potential off-target actions of PST3093, its interaction with a panel of 50 ligands, potentially relevant to off-target effects, was carried out by Eurofins (Taiwan) on crude membrane preparations according to Eurofins described procedures. PST3093 was tested at the concentration of 10 μM.

#### Functional measurements in isolated myocytes

To ensure stabilization of drug effect, effects on functional parameters were analyzed after incubating cells with PST3093 or vehicle (control) for at least 30 min. Difference between means was thus tested by group comparison. All experiments were performed at 35 °C.

### Na^+^/K^+^ ATPase current (I_NaK_)

I_NaK_ was recorded in isolated rat left ventricular (LV) myocytes as the holding current at −40 mV under conditions enhancing I_NaK_ and minimizing contamination by other conductances (Rocchetti et al., 2003; Torre et al., 2021). I_NaK_ inhibition by the compounds was expressed as percent reduction of ouabain (1 mM)-sensitive current; efficacy was expressed as the compound concentration exerting 50% inhibition (IC_50_) effect.

### Intracellular Ca^2+^ dynamics

Ca^2+^-dependent fluorescence (Fluo4-AM) was recorded in field stimulated (2 Hz) or patch-clamped rat LV myocytes and quantified by normalized units (F/F_0_). Ca^2+^ dynamics was characterized through the properties of voltage-induced Ca^2+^ transients (CaT, amplitude and decay kinetics) and caffeine-induced ones, the latter estimating total SR Ca^2+^ content (Ca_SR_). Caffeine was applied 20 seconds after the last of a CaT sequence at 2 Hz. Resting Ca^2+^ (Ca_rest_) was measured just before caffeine application to reflect the equilibrium value of cytosolic Ca^2+^.

In patch-clamped myocytes, SR Ca^2+^ uptake rate was evaluated through a “SR loading” voltage protocol, specifically devised to examine the system at multiple levels of SR Ca^2+^ loading and to rule out Na^+^/Ca^2+^ exchanger (NCX) contribution. Current through L-type Ca^2+^ channel (I_CaL_) was simultaneously recorded and the excitation–release (ER) “gain” was calculated as the ratio between CaT amplitude and Ca^2+^ influx through I_CaL_ up to CaT peak (Rocchetti et al., 2005) (protocol in Figure S1).

### Electrical activity

Action potentials (APs) were recorded (I-clamp) from isolated guinea-pig LV myocytes, selected because of the AP similarity to the human one (Odening et al., 2021), under Tyrode superfusion. AP duration at 50% and 90% repolarization (APD_50_ and APD_90_) and diastolic potential (E_diast_) were measured 1) during steady state pacing at several rates; 2) dynamically upon stepping between two rates (APD_90_ adaptation). During steady state pacing, short-term APD_90_ variability (STV) was calculated from 20-30 subsequent APD_90_ values according to Eq. 1 (Altomare et al., 2015):

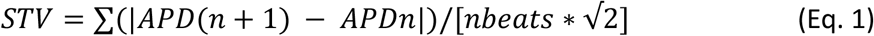

The kinetics of APD_90_ adaptation was quantified by the time constant (τ) of exponential data fitting.

#### In-vivo studies

##### Acute drug toxicity in mice

Acute toxicity of PST3093 was preliminarily evaluated in male Albino Swiss CD-1 mice by identifying the dose causing 50% mortality (LD_50_, mg.kg^-1^ body weight) at 24 hours after i.v. injection. PST3093 was dissolved in DMSO and injected at 50, 100, 200, and 250 mg kg^-1^ (4 animals for each group). Higher PST3093 dose levels were not tested due to the solubility limit of the compound; control animals received the vehicle only (DMSO).

Acute toxicity of istaroxime in male CD1 mice was also evaluated for comparison, by testing i.v. dose levels of 15, 22, 27 and 33 mg kg^-1^ (5 animals for each group). Control animals received the vehicle only (saline).

##### Hemodynamic studies in rats with diabetic cardiomyopathy

Healthy and STZ rats were studied by transthoracic echocardiographic under urethane anaesthesia (1.25 g.kg^-1^ i.p.). Left-ventricular end-diastolic (LVEDD) and end-systolic (LVESD) diameter, posterior wall (PWT) and interventricular septal (IVST) thickness were measured according to the American Society of Echocardiography guidelines (Lang et al., 2006). Fractional shortening was calculated as FS = (LVEDD-LVESD)*LVEDD^-1^. Trans-mitral flow velocity was measured (pulsed Doppler) to obtain early and late filling velocities (E, A waves) and E wave deceleration time (DT). DT was also normalized to E wave amplitude (DT/E ratio). Peak myocardial systolic (s’) and diastolic velocities (e’ and a’), were measured at the mitral annulus by Tissue Doppler Imaging (TDI). Two-dimensional LV mass and its relative index to body weight, were estimated in healthy and STZ rats.

According to our previous study (Torre et al., 2021), PST3093 was i.v. infused at 0.22 mg.kg^-1^ (0.16 ml min^-1^); echocardiographic parameters were measured before and at 15 and 30 min during the infusion. Istaroxime (0.22 mg.kg^-1^) and digoxin (0.11 mg.kg^-1^), both infused for 15 min, were used as comparators. Previous studies demonstrated that neither urethane anaesthesia, nor the infusion regimen per se affected echocardiographic parameters (Torre et al., 2021).

##### Statistical analysis

Individual means were compared by paired or unpaired *t*-test; multiple means were compared by one or two-way ANOVA for repeated measurements (RM) plus post-hoc Tukey’s multiple comparisons; drug-induced changes in overall curve steepness were defined according to significance of the “factor X group” interaction. Data are reported as mean ± SEM; p<0.05 defined statistical significance of differences in all comparisons. Number of animals (N) and/or cells (n) are shown in each figure legend.

##### Chemicals

Istaroxime {PST2744: [E,Z]-3-[(2-aminoethoxy)imino]-androstane-6,17-dione hydrochloride} and its metabolite PST3093 {(E,Z)-[(6-beta-hydroxy-17-oxoandrostan-3-ylidene)amino]oxyacetic acid} were synthesized and produced at Prassis Research Institute and Sigma-Tau Pharmaceutical Company and then by CVie Therapeutics Limited and WindTree Therapeutics. The batch of PST3093 utilized for *in-vitro* and *in-vivo* studies was a 1:1 mixture of oxime E:Z isomers. It was synthesized according to standard procedures, characterized by NMR spectroscopy and its purity (about 95%) was assessed by HPLC (Figure S2). Digoxin was purchased from Sigma-Aldrich. Istaroxime (PubChem CID: 9841834); digoxin (PubChem CID: 2724385).

## Results

### Chemical structure of PST3093

PST3093 is the final metabolite of istaroxime (Gheorghiade et al., 2008); its chemical structure is shown in Figure 1A. Compared to istaroxime, PST3093 retains the oxime moiety at position 3 with the amino-chain oxidized into a carboxylic chain, while the 6-keto group of istaroxime is stereo selectively reduced to a 6β-hydroxyl group.

**Figure 1.**
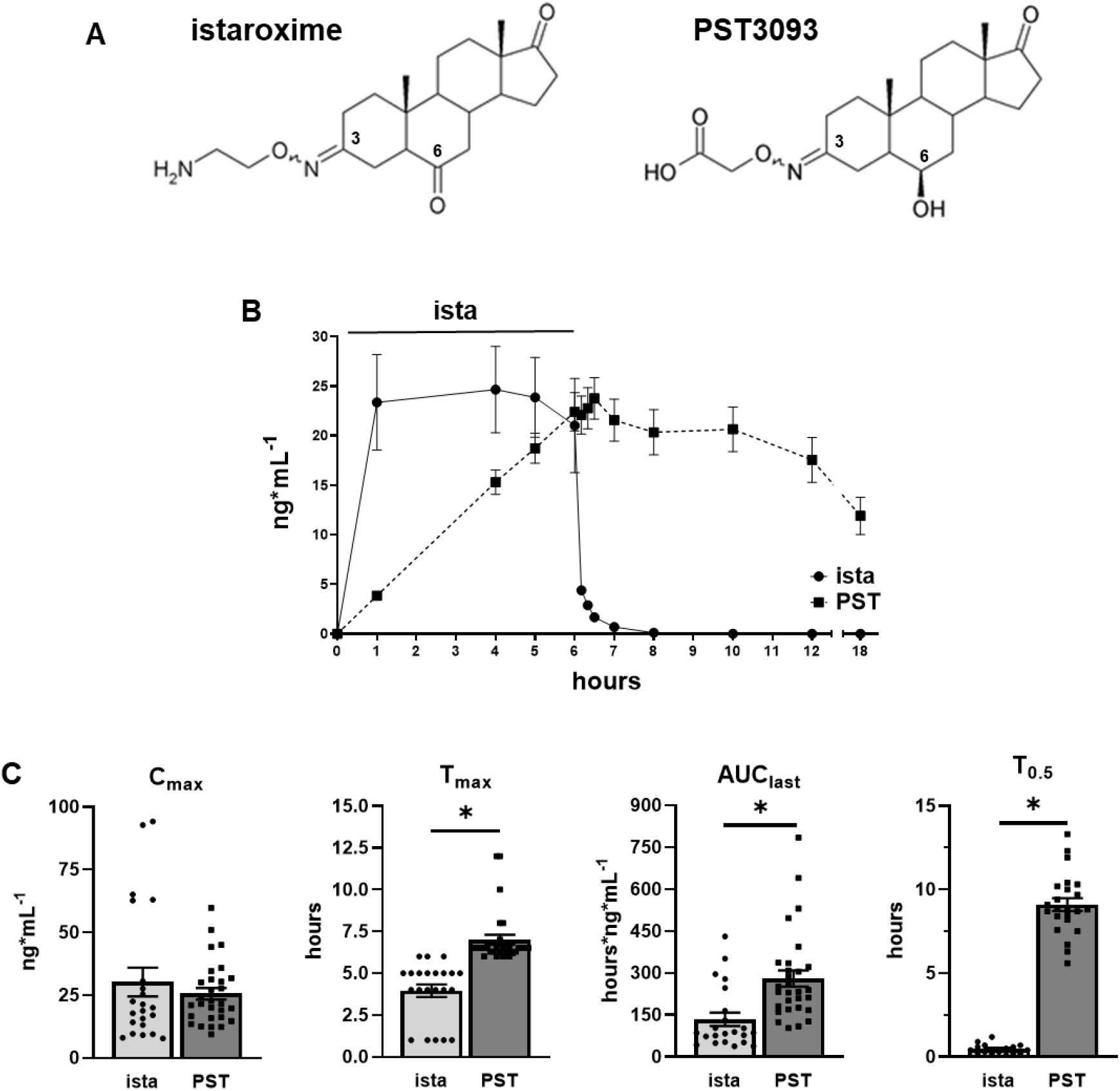
Chemical structure of istaroxime and its metabolite PST3093 (A) and PKs in humans (B-C). **A)** The main metabolic pathways of istaroxime are reduction of the carbonyl in position 6 catalyzed by carbonyl reductases and oxidative deamination of the primary amino group catalyzed by MAOs or tissue-bound semicarbazide-sensitive amine oxidase. **B)** Istaroxime (N=22) and PST3093 (N=29) plasma levels evaluated over 6 hours istaroxime infusion in HF patients at 1 μg/kg/min and up to 12 hours after wash-out (from HORIZON-HF study, NCT00616161). **C)** Statistics of the maximal observed concentration (C_max_), the time of C_max_ (T_max_), the area under the curve from the start of drug administration to the time of final sampling (AUC_last_) and the plasma half-life (T_0.5_); *p<0.05 vs istaroxime (unpaired *t*-test). Data are the mean ± SEM.

### Pharmacokinetics (PKs)

Figure 1B shows istaroxime and PST3093 plasma levels over a 6-hours istaroxime infusion at 1 μg kg^-1^ min^-1^ and up to 12 hours after discontinuation of infusion. All patients (Gheorghiade et al., 2008) showed measurable istaroxime levels until 10 min after stopping the infusion, drug levels decreased rapidly thereafter and just one patient had a quantifiable istaroxime level 2 hours after the end of the infusion. PST3093 plasma levels increased with a lag from the start of istaroxime infusion (as expected for a metabolite) being detectable in all patients from 1 hour after the start of infusion. Plasma PST3093 levels remained detectable long after discontinuation of the infusion, up to the last sample at 12 hours after wash out (Figure 1C). The data suggest that, if istaroxime infusion had continued beyond 6 hours, the metabolite would have accumulated further.

In quantitative terms, PST3093 had a plasma half-life (T_0.5_) of about 9 hours, i.e. substantially longer than that of istaroxime (less than 1 hour), leading to a huge enhancement of the AUC_last_ index for PST3093; while the maximal observed concentration (C_max_) of PST3093 was similar to istaroxime one at this infusion rate, the time of its observation (T_max_) was longer for PST3093 in comparison to istaroxime (Figure 1C).

### Effect of PST3093 on Na^+^/K^+^ ATPase

Compounds effects on Na^+^/K^+^ ATPase activity were tested in a range of concentrations from 10^−9^ to 10^−4^ M (Figure 2A). Na^+^/K^+^ ATPase from dog kidney had a specific baseline activity of 14 μmol.min^-1.^mg^-1^ protein. The reference compound istaroxime inhibited Na^+^/K^+^ ATPase activity with IC_50_ of 0.14 ± 0.02 µM in dog kidney (Figure 2A), which corresponds to a higher affinity as compared to that observed in rat renal preparations (IC_50_ of 55 ± 19 μM from (Torre et al., 2021)). PST3093 did not inhibit the Na^+^/K^+^ ATPase activity up to 100 µM, the maximal tested concentration (Figure 2A).

**Figure 2.**
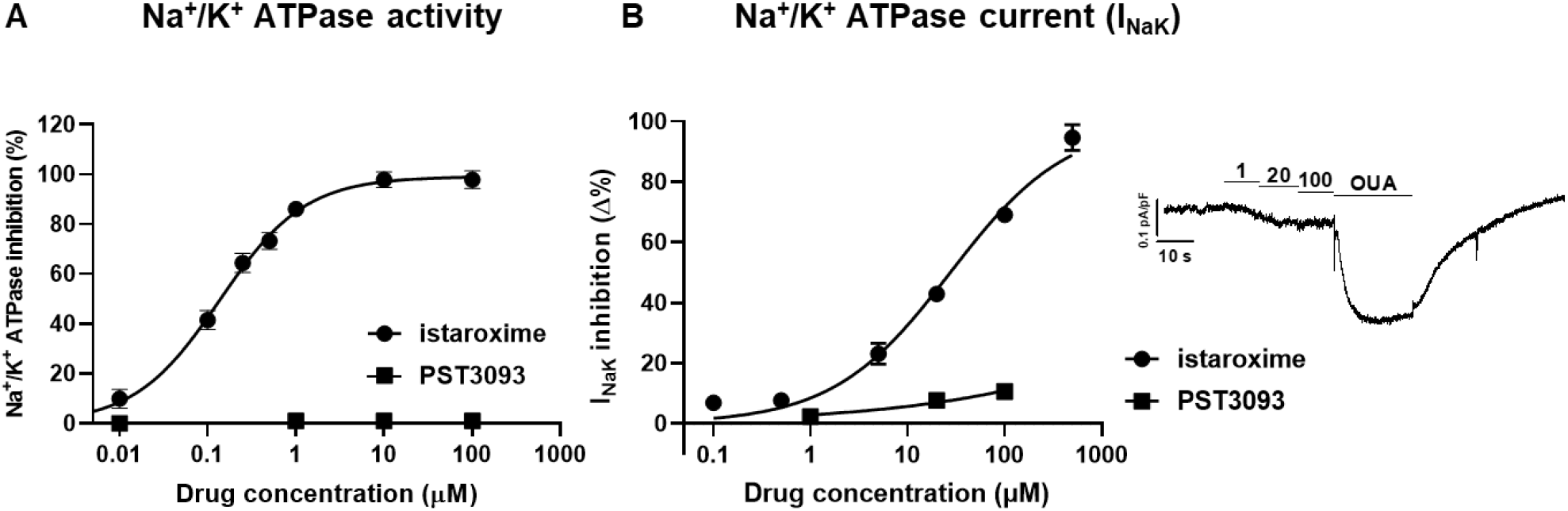
Modulation of Na^+^/K^+^ ATPase activity. **A)** inhibition of Na^+^/K^+^ ATPase activity by istaroxime and PST3093 in dog renal preparations (N=3); **B)** concentration-response curves for I_NaK_ inhibition by PST3093 (n=15) and istaroxime (modified from (Torre et al., 2021)) in rat LV myocytes; I_NaK_ recording under increasing concentrations of PST3093 and finally to ouabain (OUA as reference) is shown on the right. Data are the mean ± SEM.

PST3093 effects on Na^+^/K^+^ ATPase current (I_NaK_) were furtherly evaluated in rat LV myocytes in comparison to istaroxime (Figure 2B). The estimated IC_50_ for I_NaK_ inhibition by istaroxime was 32 ± 4 µM, (from (Torre et al., 2021)); for PST3093 a barely detectable inhibition (9.2 ± 1.1%) was observed at the limit concentration for solubility (100 µM), an effect that can be considered insignificant.

### Effect of PST3093 on SERCA ATPase activity

#### Effects on SERCA2a activity in normal and diseased myocardial preparations

In STZ rat preparations (N=30), baseline SERCA2a V_max_ was lower (by −27%) than in healthy ones (N=29) (0.199±0.01 vs 0.272±0.01 μmol*min^-1^*mg^-1^ protein, p<0.05), with no difference in K_d_Ca (448±35 vs 393±22 nM, NS), similarly to what reported recently in the same setting (Torre et al., 2021). As also reported previously (Ferrandi et al., 2013), the response of enzyme kinetics parameters to modulation was species-specific: whereas in rat preparations both PST3093 and istaroxime (Figure 3) increased V_max_, in guinea-pig ones the compounds decreased K_d_Ca instead (Table S1).

**Figure 3.**
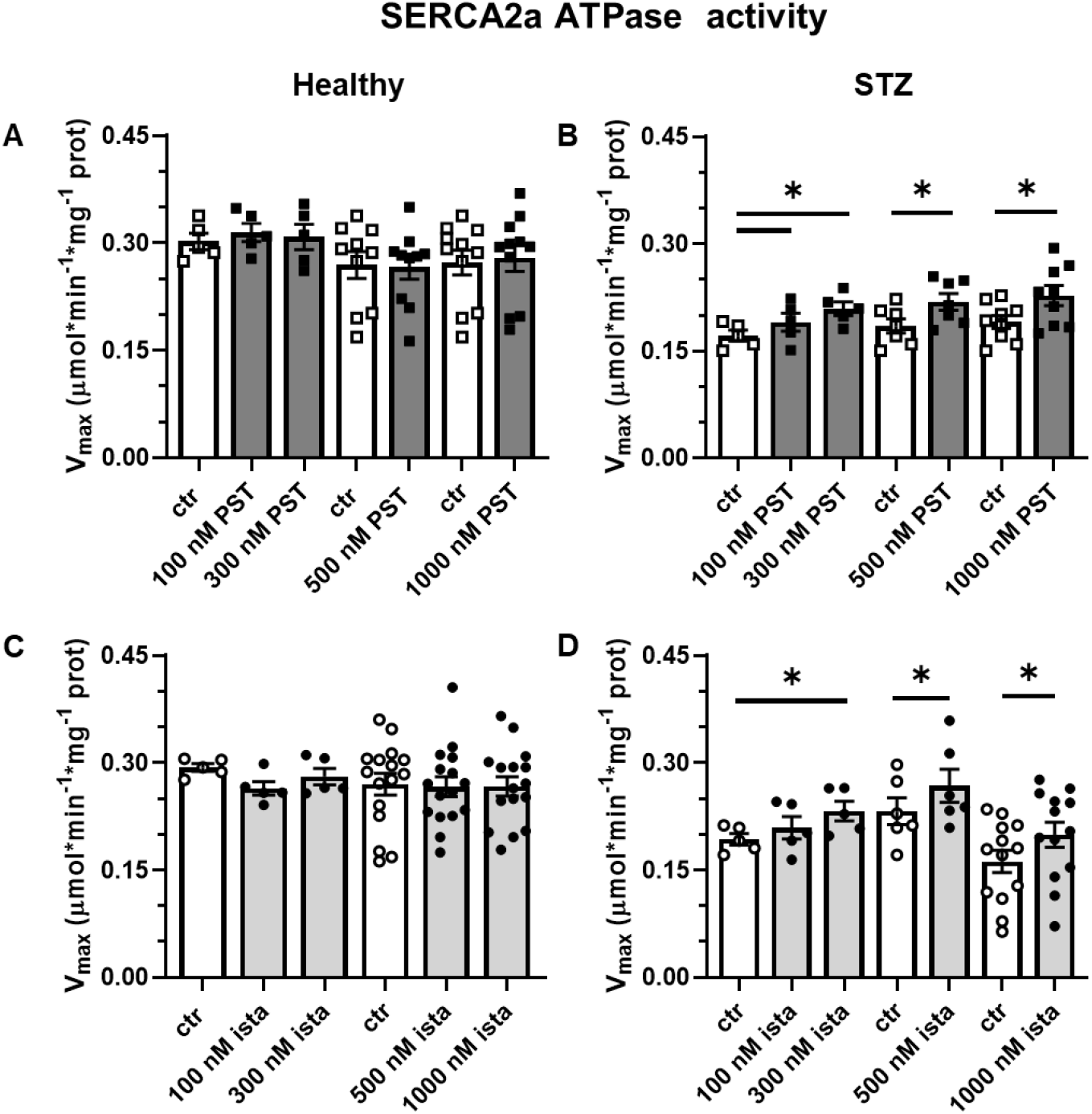
Modulation of SERCA2a ATPase activity in healthy and diseased (STZ) preparations. Effect of PST3093 (N=5-11) **(A-B)** and istaroxime (N=5-16) **(C-D)** on SERCA2a V_max_ estimated from Ca^2+^-dose response curves in cardiac homogenates from healthy and diabetic (STZ) rats. Internal controls (ctr) are provided. Data are the mean ± SEM. * p<0.05 vs ctr (RM one-way ANOVA plus post-hoc Tukey’s multiple comparisons test or paired t-test).

Over the whole range of concentration tested (0.1 - 1µM), PST3093 and istaroxime failed to affect ATPase Ca^2+^-dependency in healthy rat preparations (Figure 3A,C), but similarly increased V_max_ in STZ ones (e.g. +22% and +20%, respectively at 300 nM) with thresholds at 100 nM and 300 nM for PST3093 and istaroxime, respectively (Figure 3B,D); SERCA2a K_d_Ca in rat preparations was affected by neither istaroxime nor PST3093. Thus, PST3093 and istaroxime displayed similar potency in ameliorating disease-induced depression of SERCA2a ATPase activity.

PST3093 and istaroxime effect was also detected in preparations from normal guinea-pig hearts, where they both reduced the SERCA2a K_d_Ca value by about 20% at 100 nM (Table S1).

To summarize, PST3093 and istaroxime equally stimulated SERCA2a ATPase activity in preparations including PLN. Regardless of the kinetic parameter affected, a stimulatory effect was present in healthy guinea-pig microsomes and in rat homogenates from the STZ disease model.

#### Dependency of SERCA stimulation by PST3093 on PLN

In a range of concentrations from 30 to 1000 nM, PST3093 and istaroxime failed to affect SERCA1 activity in the absence of PLN in skeletal muscle preparations (Figure 4A,C). Reconstitution with the PLN_1-32_ fragment markedly reduced SERCA1 affinity for Ca^2+^ (K_d_Ca increased by 23-26%) (Figure 4B,D). Under this condition, both PST3093 (Figure 4B) and istaroxime (Figure 4D) dose-dependently reversed PLN-induced shift in K_d_Ca with an EC_50_ of 39 nM and 40 nM respectively.

**Figure 4.**
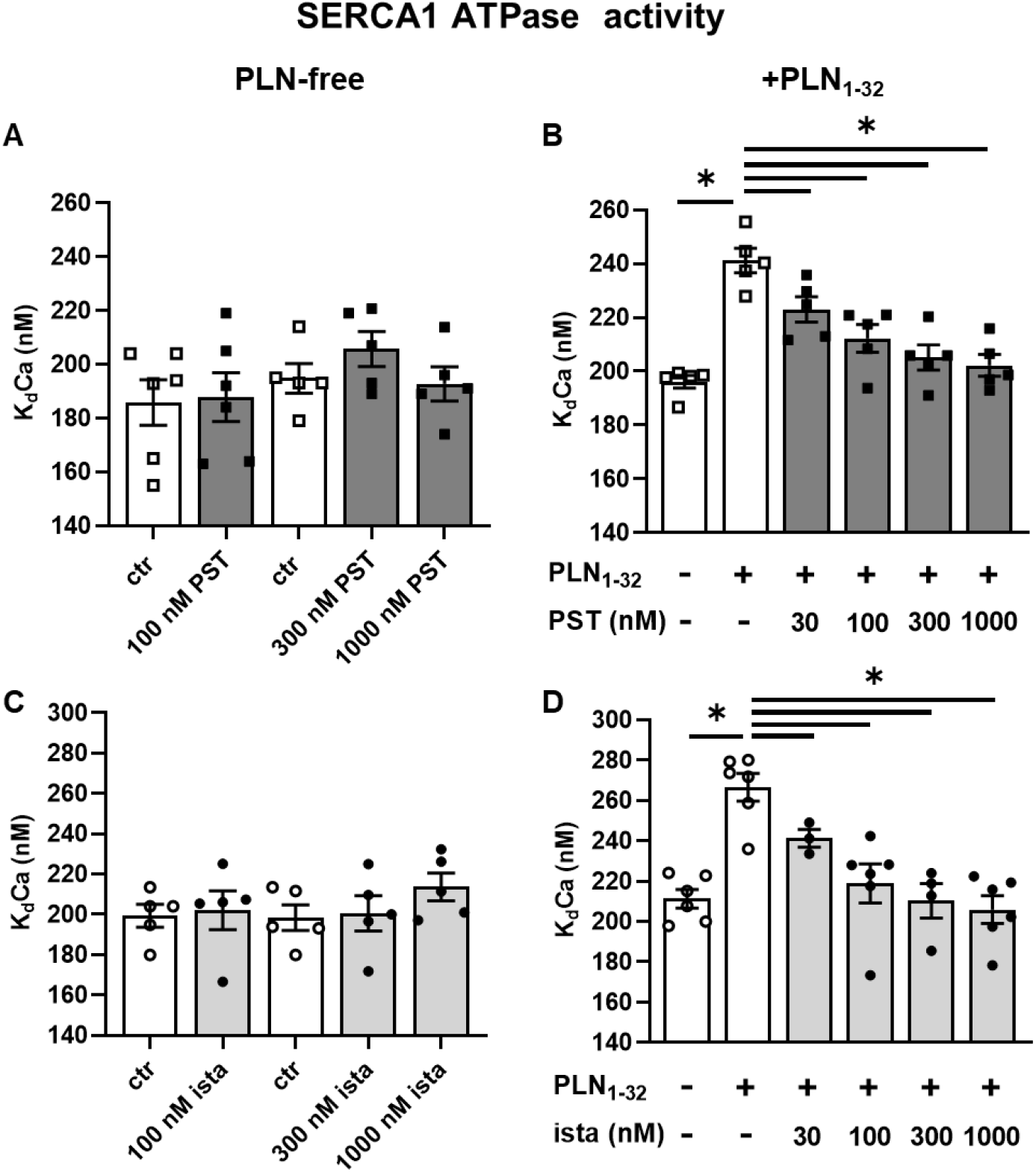
Modulation of SERCA1 ATPase activity. Concentration-dependency of PST3093 and istaroxime modulation of SERCA1 K_d_Ca in guinea-pig skeletal muscle microsomes containing SERCA1 alone **(A, C)** (N=5,6) and after reconstitution with the PLN_1-32_ fragment **(B, D)** (N=5,6). Data are the mean ± SEM. *p<0.05 (RM or mixed model one-way ANOVA plus post-hoc Tukey’s multiple comparisons).

##### PST3093 interaction with targets other than SERCA

The targets panel (50 items) included membrane receptors, key enzymes, ion channels and transporters, relevant to potential off-target cardiac and extra cardiac effects (list in Table S2); PST3093 was tested at the concentration of 10 μM. None among the 50 items met criteria for significance of interaction. Thus, at least for the ligands shown in Table S2, no off-target action of PST3093 is expected.

##### Effects of PST3093 on intracellular Ca^2+^ dynamics in cardiac myocytes

###### Effect on intracellular Ca^2+^ dynamics under field-stimulation

Ca^2+^ dynamics were analysed in field stimulated (2 Hz, Figure 5) rat LV myocytes isolated from healthy or STZ rats.

**Figure 5.**
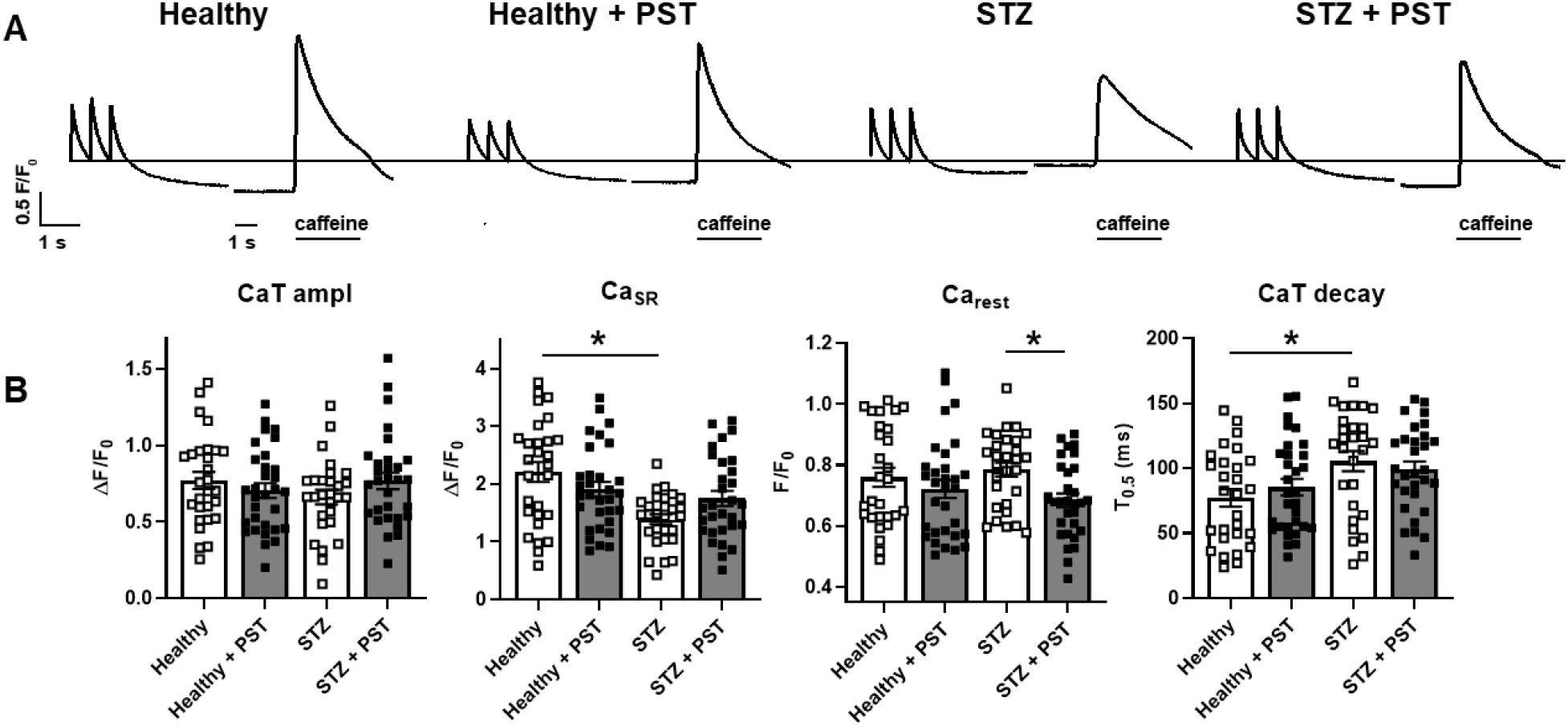
Modulation of intracellular Ca^2+^ handling in field stimulated myocytes from healthy and diseased (STZ) hearts. **A)** Representative recordings of Ca^2+^ transients (CaT) triggered by steady-state electrical stimulation at 2Hz in intact cells, followed after 20 secs by caffeine-induced Ca^2+^ release measuring SR Ca^2+^ content (Ca_SR_). PST3093 (1 µM) was tested in normal (healthy) and diseased (STZ) myocytes. **B)** Statistics for CaT amplitude, Ca_SR_, Ca^2+^ at pause end (Ca_rest_), time for 50% CaT decay (T_0.5_). CTR N=3 (n=28 w/o PST3093, n=31 with PST3093), STZ N=4 (n=28 w/o PST3093, n=30 with PST3093). *p<0.05 (one-way ANOVA plus post-hoc Tukey’s multiple comparisons).

STZ myocytes had a lower SR Ca^2+^ content (Ca_SR_) and a slower CaT decay than healthy ones; however, CaT amplitude remained unchanged. These changes are compatible with reduced SERCA2a function, possibly compensated by APD prolongation, known to increase cell Ca^2+^ content (Torre et al., 2021). Whereas in healthy myocytes 1 µM PST3093 failed to affect any of the Ca^2+^ dynamics parameters, in STZ-myocytes PST3093 reduced the quiescence Ca^2+^ (Ca_rest_) prior to caffeine application and partially restored Ca_SR_ and CaT decay. Moreover, PST3093 restored in STZ-myocytes the distribution of Ca_SR_ values peculiar of healthy ones. Comparable results have been obtained with istaroxime at a concentration marginally affecting Na^+^/K^+^ ATPase (Torre et al., 2021). Taken together, this observation suggests that PST3093 improved Ca^2+^ sequestration into the SR during the post-train quiescence period.

###### Effect on SR Ca^2+^ uptake function in V-clamped myocytes

STZ-induced changes in repolarization affect Ca^2+^ handling in a direction masking SERCA2a downregulation (Torre et al., 2021). Thus, the “SR loading” protocol (see methods and Figure S1) was performed under V-clamp and used to assess SR Ca^2+^ uptake under conditions emphasizing SERCA2a role. In STZ myocytes, as compared to healthy ones, SR reloading was significantly depressed (both in terms of CaT amplitude and ER gain) and CaT decay was slower at all time-points during reloading (Figure 6A). These changes are compatible with depressed SERCA2a function (Torre et al., 2021). PST3093 (1 µM) was tested in STZ myocytes (Figure 6B), where it sharply accelerated CaT decay, restoring the profile observed in healthy myocytes. Albeit less evident, drug-induced changes in CaT and ER gain pointed in the same direction. Comparable results have been obtained with istaroxime at a concentration marginally affecting Na^+^/K^+^ ATPase (Torre et al., 2021).

**Figure 6.**
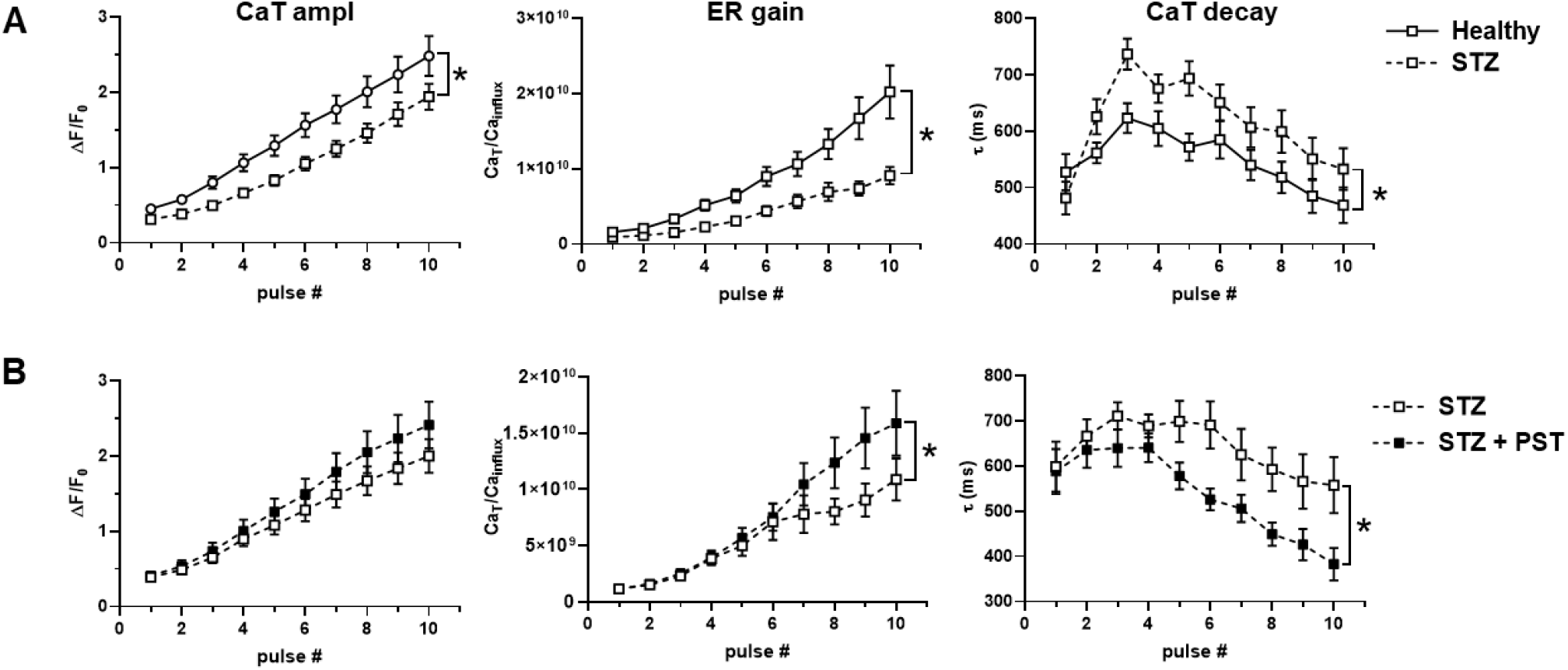
Modulation of SR Ca^2+^ uptake under NCX inhibition in V-clamped myocytes from STZ hearts. **A)** Disease (STZ) effect on SR Ca^2+^ loading in patch-clamped myocytes. SR Ca^2+^ loading by a train of V-clamp pulses was initiated after caffeine-induced SR depletion; NCX was blocked by Na^+^ substitution to identify SERCA2a-specific effects (see Methods and Figure S1); myocytes from healthy (N=9, n=32) and diseased hearts (STZ, N=6, n=31) are compared. **B)** PST3093 effect in STZ myocytes (N=4, w/o PST3093 n=18, with PST3093, n=19). Panels from left to right: CaT amplitude, Excitation-Release (ER) gain (the ratio between CaT amplitude and Ca^2+^ influx through I_CaL_), time constant (τ) of CaT decay. *p<0.05 for the “interaction factor” in RM two-way ANOVA, indicating a different steepness of curves.

Overall, PST3093 restored SR function in diseased myocytes, most likely through SERCA2a enhancement, within the context of an intact cellular environment.

##### Effects of PST3093 on cellular electrical activity

To assess the electrophysiological safety of PST3093, its effects on AP of LV myocytes were investigated. Guinea-pig myocytes were used, instead of rat ones, because their AP is closer to the human one.

PST3093 (100 nM) marginally reduced APD_50_ at all pacing rates, leaving the other AP parameters unchanged (Figure 7A-B). Notably, also APD rate-dependency at steady-state and the kinetics of APD adaptation following a step change in rate, were unaffected by the agent (Figure 7C). STV of APD_90_, a reporter of repolarization stability, was also unaffected by PST3093 at all pacing rates (Figure 7D-E). Except for the absence of APD_50_ reduction, similar results were obtained with PST3093 at 1 µM (Figure S3).

**Figure 7.**
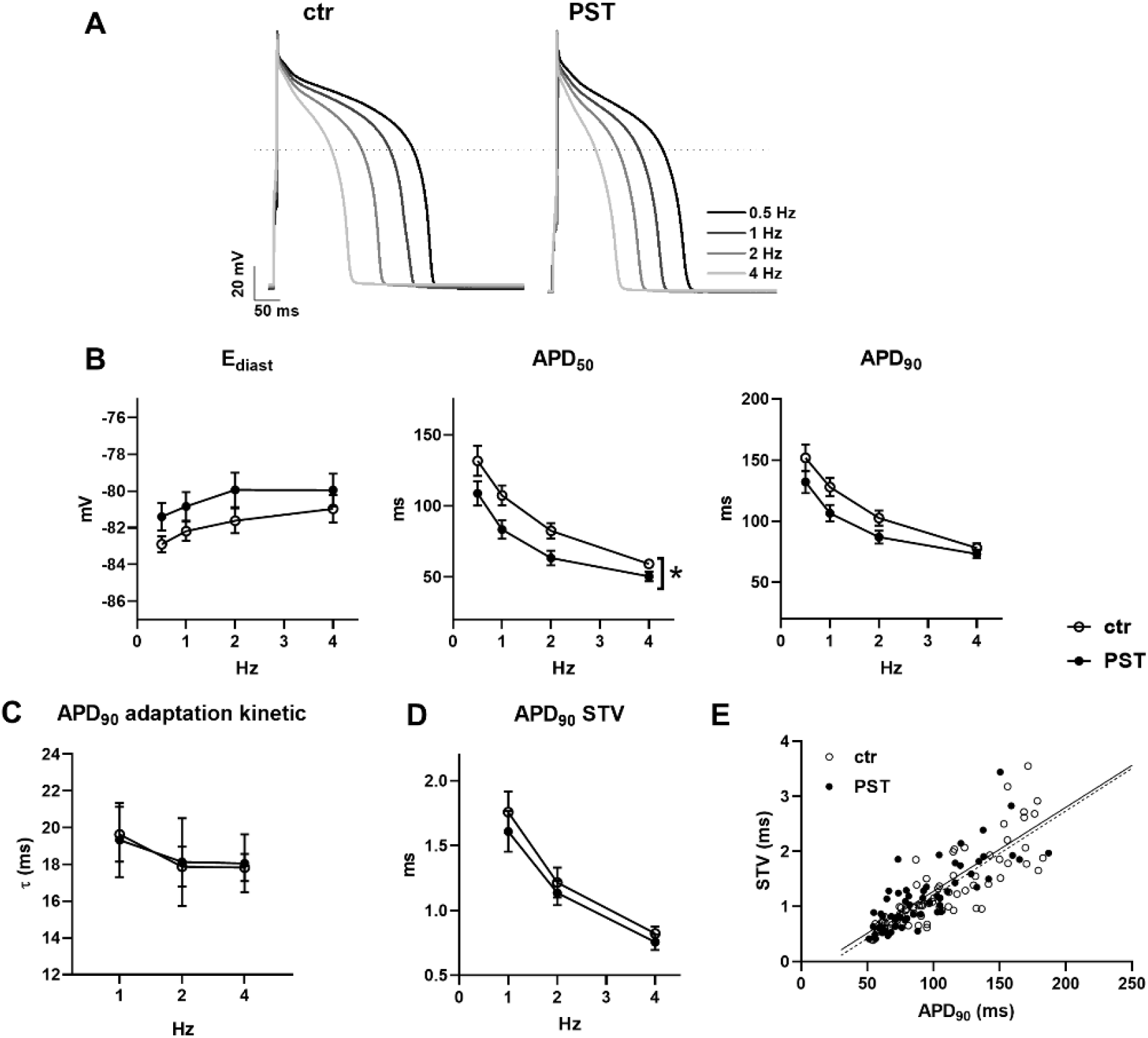
Modulation of electrical activity in guinea pig myocytes. The effect of 100 nM PST3093 was tested on action potential (AP) parameters and their steady-state rate-dependency in guinea pig myocytes (N=5). **A)** representative APs recorded at 0.5, 1, 2, 4 Hz in control (left) and with 100 nM PST3093 (right). **B)** Effect on the rate dependency of diastolic potential (E_diast_) and AP duration (APD_50_, APD_90_) (n≥24 w/o PST3093, n≥22 with PST3093). **C)** Effect on the time constant (τ) of APD_90_ adaptation following a step change in rate (n≥22 w/o PST3093, n≥16 with PST3093). **D)** Effect on the rate-dependency of APD_90_ short term variability (STV) (n≥24 w/o PST3093, n≥20 with PST3093). **E)** Effect on the correlation between STV of APD_90_ and APD_90_ values; data from 1, 2, 4 Hz were pooled. *p<0.05 for the “interaction factor” of RM two-way ANOVA. The effect of PST3093 at a higher concentration (1 µM) is reported in Figure S3.

The paucity of PST3093 effects on the AP is consistent with the absence of hits in the analysis of PST3093 interaction (up to 10 μM) with molecular targets other than SERCA2a, among them ion channels and transporters (Table S2).

##### In vivo acute toxicity in mice

*In vivo* toxicity after i.v. injection was investigated in mice for PST3093 and istaroxime. PST3093 was well tolerated and did not cause death up to 250 mg/kg. The LD_50_ was not calculated since no deaths occurred at the maximal usable dose (limited by solubility). The LD_50_ for istaroxime was 23.06 mg/kg. The main signs of toxicity were prostration, gasping and convulsions. In most of the animals, death occurred within 5 min after istaroxime administration. Post-mortem examination revealed pulmonary oedema and/or haemorrhages and generalized organ congestion. No remarkable alterations were found in the surviving animals.

Therefore, when i.v. administered, PST3093 was far less toxic than istaroxime, a result ascribable to its lack of effects on the Na^+^/K^+^ ATPase.

##### Modulation of cardiac function in vivo in rats with diabetic cardiomyopathy

###### Features of the disease model

Fasting hyperglycaemia, polydipsia, polyuria and polyphagia ensued 1 week after STZ injection; none of these symptoms was observed in healthy rats. Eight weeks after STZ, total body weight (BW) was substantially lower in STZ rats; LV mass (by echo) was reduced in absolute value but, when normalized to BW, was not significantly different in STZ rats (Table S3). The STZ model was completely characterized in a previous work of ours (Torre et al., 2021).

In this study a comprehensive echocardiographic analysis of STZ rats in comparison to healthy ones was performed. Figure 8 compares some echo indexes in STZ vs healthy rats; Table S4 lists all the measured echo parameters in the two groups. Heart rate (HR) was lower in STZ rats (−20%; p<0.05) and stroke volume (SV) unchanged; nonetheless, differences in cardiac output (CO) between the two groups did not achieve significance. *Systolic indexes:* in STZ rats LV end-systolic diameter (LVESD) was larger and the ejection fraction (EF) reduced; fractional shortening (FS) and systolic tissue velocity (s’) were depressed. *Diastolic indexes:* in STZ rats LV end-diastolic diameter (LVEDD) tended to be larger; peak E wave velocity (E) was slightly smaller and A wave velocity (A) was unchanged (E/A unchanged); E wave deceleration time (DT) was unchanged in absolute, but the DT/E ratio increased. Changes in early (e’) and late (a’) diastolic tissue velocities paralleled those in E and A waves; therefore, the e’/a’ and E/e’ ratios did not differ between STZ and healthy rats.

**Figure 8.**
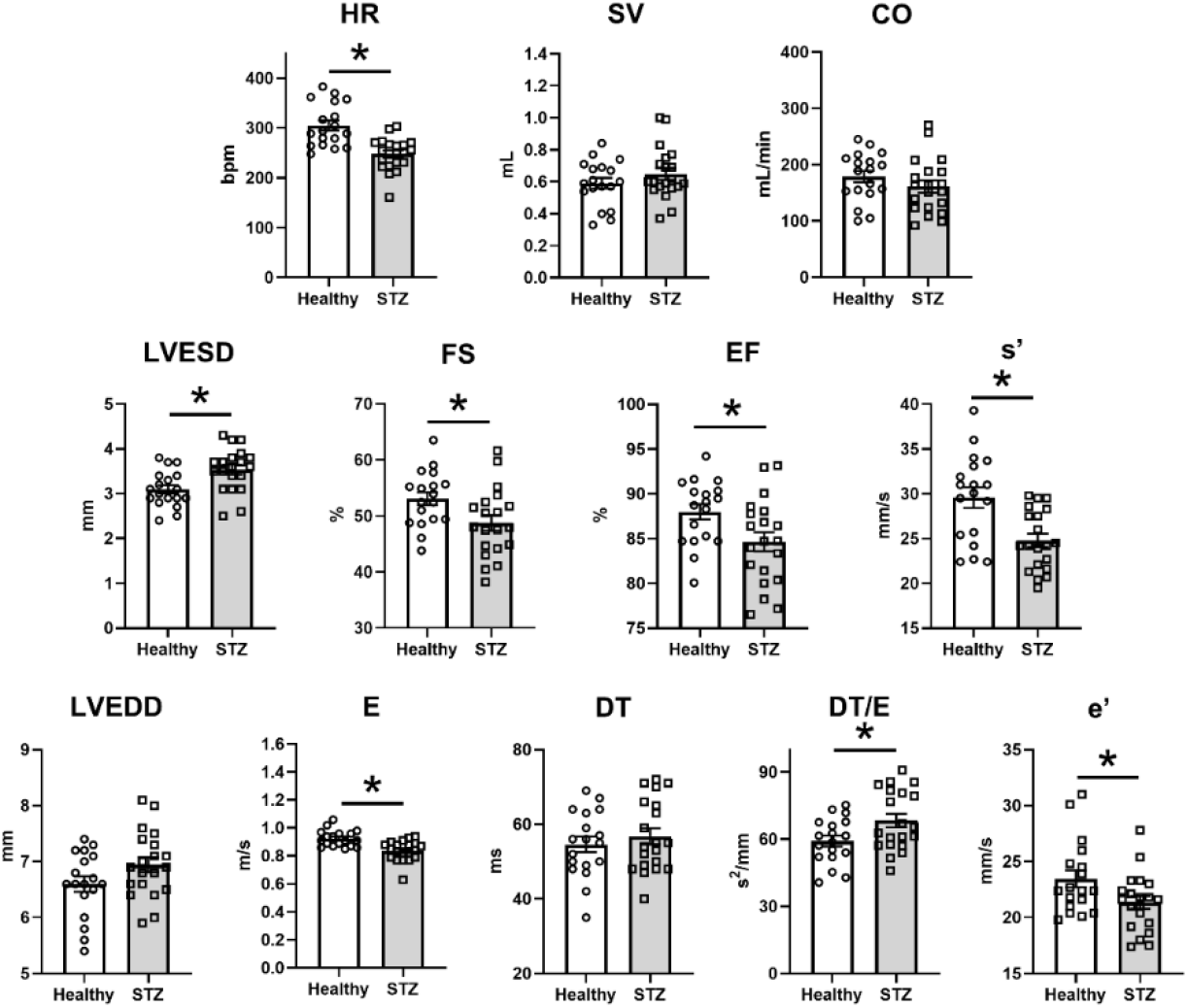
Disease (STZ) effect on *in-vivo* echocardiographic parameters. Echocardiographic parameters are compared between healthy rats (N=18) and 8 weeks after STZ treatment (N=20). Top row, global function parameters: HR: heart rate; SV: stroke volume; CO: cardiac output. Mid row, systolic function parameters: LVESD: left ventricular end-systolic diameter; FS: fractional shortening; EF: ejection fraction; s’: peak systolic tissue velocity. Bottom row, diastolic function parameters: LVEDD: left ventricular end-diastolic diameter; E: E wave amplitude; DT: E wave deceleration time; DT/E: normalized DT; e’: peak early diastolic tissue velocity. *p<0.05 (unpaired *t*-test).

To summarize, in STZ rat echocardiographic abnormalities were rather subtle; nonetheless, 11 out of 21 indexes were significantly affected indicating derangements in both systolic and diastolic function.

###### Drug effects in the disease model

The *in vivo* acute effect of PST3093 (0.22 mg.kg^-1.^min^-1^) on echo indexes of STZ rats was investigated at 15 and 30 minutes of infusion (Figure 9). Data at 15 min were also obtained with istaroxime (0.22 mg.kg^-1.^min^-1^) and digoxin (0.11 mg.kg^-1.^min^-1^) (Table 1).

**Table 1.** Effects of PST3093 (0.22 mg.kg^-1^.min^-1^), istaroxime (0.22 mg.kg^-1^.min^-1^) and digoxin (0.11 mg.kg^-1^.min^-1^) on echo indexes of STZ rats at 15 minutes of infusion. Data are the mean ± SEM. #p<0.05 vs before drug infusion (paired *t*-test).

| Echo parameters |  | Istaroxime |  | PST3093 |  | Digoxin |  |
| --- | --- | --- | --- | --- | --- | --- | --- |
|  |  | before | @ 15 min | before | @15 min | before | @ 15 min |
| Morphometric parameters | IVSTd (mm) | 1,91±0,09 | 1,97±0,16 | 1,89±0,09 | 1,86±0,11 | 1,79±0,07 | 1,81±0,04 |
|  | PWTd (mm) | 1,6±0,07 | 1,68±0,09 | 1,46±0,05 | 1,47±0,07 | 1,71±0,08 | 1,66±0,1 |
|  | LVEDD (mm) | 7,25±0,15 | 7,1±0,2 | 7,22±0,17 | 7,57±0,17 | 6,76±0,15 | 6,67±0,15 |
|  | IVSTs (mm) | 2,4±0,1 | 2,45±0,12 | 2,16±0,11 | 2,31±0,13 | 2,25±0,14 | 2,31±0,1 |
|  | PWTs (mm) | 2,33±0,16 | 2,42±0,15 | 2,43±0,12 | 2,62±0,14 | 2,47±0,16 | 2,68±0,14 |
|  | LVESD (mm) | 3,82±0,21 | 3,8±0,2 | 3,89±0,18 | 3,66±0,15 | 3,53±0,17 | 3,14±0,14 # |
| Systolic parameters | FS (%) | 47,21±2,4 | 46,72±2,1 | 46,2±2,02 | 51,8±1,89 # | 47,62±1,9 | 53,09±1,8 # |
|  | s' (mm/s) | 23,37±0,9 | 23,05±0,7 | 22,0±0,72 | 23,4±0,5 # | 23,5±0,9 | 26±1,5 # |
|  | EF (%) | 83 ± 2 | 82 ± 62 | 82 ± 1,9 | 87 ± 1,5 # | 84±1,5 | 88±1,3 # |
| Diastolic parameters | E (mm/s) | 0,91±0,04 | 0,94±0,04 | 0,91±0,06 | 1,05±0,07 # | 0,831±0,03 | 0,924±0,06 |
|  | A (mm/s) | 0,66±0,04 | 0,78±0,05 # | 0,79±0,08 | 0,87±0,07 | 0,59±0,05 | 0,79±0,05 # |
|  | E/A | 1,39±0,06 | 1,22±0,08 | 1,18±0,05 | 1,22±0,04 | 1,48±0,11 | 1,2±0,06 # |
|  | DT (ms) | 46,62±4,1 | 34,12±2,5 # | 48,5±3,8 | 40,2±3,0 | 53,4±3,8 | 44,1±3,2 # |
|  | DT/E | 51,91±4,9 | 36,66±3,3 # | 56,7±6,5 | 40,39±4,8 # | 65,21±5,3 | 50,18±5,4 # |
|  | E/DT | 21,28±3,2 | 28,98±2,8 # | 20,61±3,0 | 27,51±2,6 # | 16,53±1,7 | 22,66±3,0 |
|  | e' (mm/s) | 21,23±0,99 | 24,73±0,61# | 22,94±1,0 | 27,01±1,1 # | 20,9±0,5 | 23,8±0,9 # |
|  | a' (mm/s) | 27,61±1,9 | 31,35±1,5 # | 28,72±2,2 | 31,06±1,5 | 25,9±1,9 | 29,8±1,9 |
|  | e'/a' | 0,78±0,04 | 0,79±0,03 | 0,83±0,06 | 0,88±0,04 | 0,85±0,06 | 0,82±0,04 |
|  | E/e' | 43,6±2,6 | 38,29±1,6 # | 39,5±1,7 | 38,61±1,5 | 39,64±0,9 | 38,95±2,3 |
| Cardiac function | HR (bpm) | 266,2±9,8 | 316,2±7,5 # | 270±17 | 271±10 | 236±12 | 257±11 |
|  | SV (ml) | 0,71±0,04 | 0,67±0,05 | 0,702±0,05 | 0,847±0,06# | 0,589±0,03 | 0,601±0,04 |
|  | CO (ml/min) | 188,8±10,5 | 211,2±13,2# | 187,7±13,7 | 231,7±20,2# | 138,5±11,2 | 155,6±14,4 |
|  | N | 8 | 8 | 10 | 10 | 10 | 10 |

**Figure 9.**
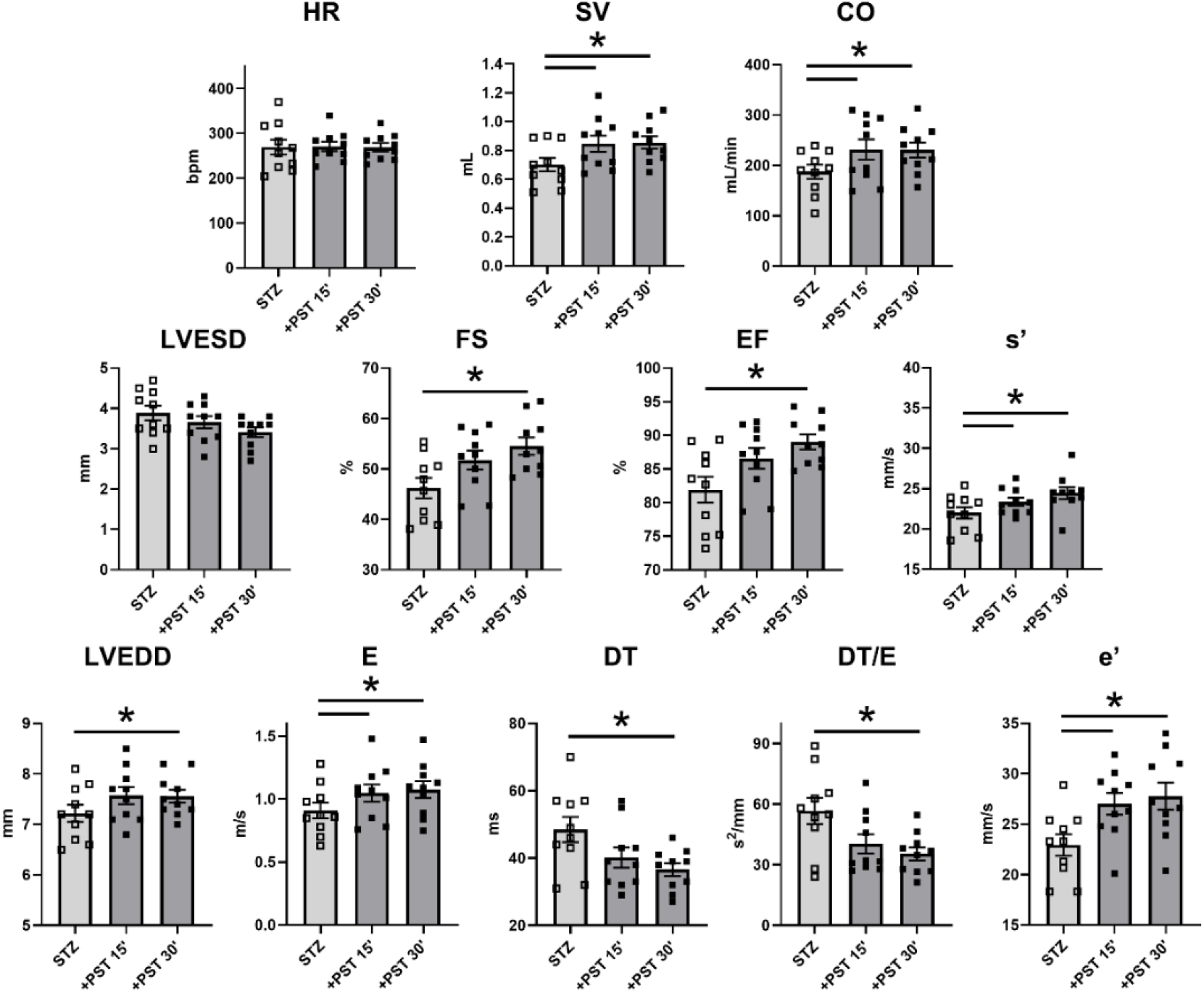
PST3093 effects on *in-vivo* echocardiographic parameters in diseased (STZ) rats. PST3093 was intravenously infused (0.22 mg.kg^-1^.min^-1^) in rats 8 weeks after STZ treatment. Echocardiographic parameters were measured before and at 15 and 30 min during drug infusion. Rows and symbols as in Figure 8; N=10 rats. Data are the mean ± SEM. *p<0.05 (RM one-way ANOVA plus post-hoc Tukey’s multiple comparisons). Effects of PST3093 on all echocardiographic parameters at 15 minutes of infusion in STZ rats in comparison to istaroxime and digoxin are reported in Table 1.

Overall, PST3093 increased SV volume and, albeit unchanged HR, it increased CO (Figure 9). *Systolic indexes*: PST3093 tended to decrease LVESD and increased FS, EF and s’. *Diastolic indexes*: PST3093 increased LVEDD, E, A and e’ (E/A and E/e’ unchanged, Table 1); DT and DT./E were reduced. PST3093 effect was almost complete at 15 min of infusion, only minor increments were observed at 30 min (Figure 9). Collectively, PST3093 improved overall cardiac function, both systolic and diastolic, beyond simple recovery of STZ-induced derangements. As shown in Figure 9, PST3093 “reversed” STZ-induced changes in 7 out of 11 indexes; moreover, five additional indexes, unaffected by STZ, were changed by PST3093 in a direction compatible with positive inotropy/lusitropy (Table 1).

PST3093 effects were only partially shared by istaroxime at the same infusion rate (Table 1). Istaroxime failed to increase SV and systolic indexes (SV, FS, EF, s’); it increased CO but, at variance with PST3093, this was because HR increased. Similar to PST3093, istaroxime shortened DT, DT/E and increased e’, a’ and A; however, it did not change E, thus reducing E/e’. Digoxin (Table 1), as expected from its inotropic effect, increased systolic indexes (FS, EF and s’); however, at variance with PST3093, it did not affect SV or CO. Notably, also digoxin improved diastolic function to some extent: E/A, DT and DT/E were reduced and e’ was increased.

## Discussion

In the present study, we have investigated effects of the istaroxime metabolite PST3093 at molecular, cellular and *in-vivo* levels. The interest in this molecule is motivated by 1) the possibility that it may actually contribute to (i.e. be endowed with) the unique mechanism of action and interesting therapeutic profile of istaroxime (inotropy and lusitropy at low proarrhythmic risk, confirmed in phase 2 clinical trials) (Carubelli et al., 2020; Gheorghiade et al., 2008; Sabbah et al., 2007; Shah et al., 2009) and 2) the possibility it would afford to test the clinical benefit associated specifically with the rescue of SERCA2a depression, which is widely recognized as the basis for many among HF abnormalities.

PST3093 effect has been tested in three experimental settings with incremental level of biological organization, including *in-vivo* measurements from diseased hearts. The consistency of effects across these three sets of experiments confers robustness to the findings.

The results of molecular studies indicate that PST3093 differs from istaroxime because it is devoid of any inhibitory activity on the Na^+^/K^+^ ATPase, while retaining SERCA2a stimulatory action. This identifies PST3093 as a “selective” SERCA2a activator. The results also indicate that, similar to istaroxime (Ferrandi et al., 2013), PST3093 may act by weakening SERCA-PLN interaction. In rat preparations, PST3093 (and istaroxime) increased SERCA2a V_max_. Albeit this apparently conflicts with the notion that interference with PLN should decrease the K_d_Ca instead (Brittsan et al., 2003), the same pattern coexisted with evidence, by several independent approaches, of istaroxime antagonism of SERCA2a-PLN interaction (Ferrandi et al., 2013). The reason for this apparent discrepancy is unclear; the observation that PST3093 effect on SR Ca^2+^ uptake was present in rat cardiac myocytes (i.e. at physiological Ca^2+^ concentrations) suggests that it may reside in specificities of the microsomal preparation.

Investigations in intact ventricular myocytes confirm a negligible effect of PST3093 on Na^+^/K^+^ pump function. Furthermore, PST3093 abolished STZ-induced intracellular Ca^2+^ abnormalities likely dependent on SERCA2a downregulation. While PST3093 clearly affected several Ca^2+^ cycling parameters under V-clamp conditions (Figure 6), its effects in field stimulated cells were apparently small (Figure 5). As we have previously shown in this experimental model (Torre et al., 2021), evaluation of Ca^2+^ handling without controlling membrane potential (field stimulation) and without disabling competing mechanisms (NCX) may be unsuitable to detect SERCA2a activation. This is why we also performed experiments under V-clamp and with disabled NCX function, shown in Figure 6. Field-stimulation experiments are nonetheless informative because they better represent drug effects at the cell level under “physiological” conditions. In particular, the reduction in Ca_rest_ indicates that PST3093 allows the other Ca^2+^ cycling parameters to be preserved (or even showing a trend to improve) at a lower cytosolic Ca^2+^ level. Considering that all Ca^2+^ homeostatic mechanisms are in place in this setting, this is precisely what should be expected from pure SERCA2a activation (Alemanni et al., 2011; Zaza and Rocchetti, 2015), i.e. improved subcellular Ca^2+^ compartmentalization. That this apparently small change in myocyte physiology has an impact on *in-vivo* cardiac performance is shown by the *in-vivo* echo measurements (Figure 9).

The present *in-vivo* studies were conducted by echocardiography in a disease model characterized by impairment of SERCA2a function (Choi et al., 2002; Torre et al., 2021). In this model, PST3093 infusion improved overall cardiac performance (SV and CO); both systolic and diastolic indexes were positively affected by the agent. Mechanistic interpretation of echocardiographic indexes is often ambiguous. For instance, both DT prolongation and shortening have been associated with deterioration of diastolic function (Eren et al., 2004; Sabbah et al., 2007). These puzzling observations can be interpreted by considering the contribution to DT of opposing factors, each prevailing in a specific condition (Mossahebi et al., 2015). At any rate, whenever HF was associated with DT shortening, istaroxime (having PST3093 as a metabolite) prolonged it (Sabbah et al., 2007; Shah et al., 2009). A further difficulty may arise from the expectation that ino-lusitropy may increase atrial contraction (A amplitude) and ventricular relaxation (E amplitude) at the same time, thus conceivably making their ratio (E/A) unable to detect drug effects on diastolic function. An approach to the interpretation of drug effects, independent of mechanistic models, is to check whether the drug counters disease-induced abnormalities. In the case of PST3093, this was true for the majority of indexes (7 out of 11), the most notable exception being a small further increase in LVEDD. While increments in LVEDD are usually associated with deterioration of systolic function, PST3093 tended to decrease LVESD instead. The LVEDD increment was indeed associated with increased SV, and EF to which it likely contributed.

With the exception of guinea-pig, PST3093 efficacy on SERCA2a function in the diseased condition consistently contrasted with the lack of effect in healthy preparations (Figures 3 and 5). This suggests that SERCA2a function, while not strictly limiting in health, may become so whenever its “reserve” is diminished. This view may not clash with the clear-cut effect of PLN knock-out in healthy murine myocytes; indeed, SERCA2a modulation by PST3093 may be, albeit functionally significant, subtler than complete PLN ablation.

While failing to increase the amplitude of CaT, PST3093 improved echo indexes of systolic function. Mechanisms at two levels may account for this observation: at the intracellular level, increased compartmentation of Ca^2+^ within the SR may improve the energetic efficiency of Ca^2+^ cycling (Shannon et al., 2001); at the organ level, improved relaxation may increase preload, with its well-known impact on systolic force (Shiels and White, 2008). Indeed, normalization of diastolic function in HF patients with preserved ejection fraction, may restore cardiac output irrespective of changes in the latter (Tobushi et al., 2017). On the other hand, digoxin, whose mechanism of action is purely inotropic, accelerated early relaxation (DT shortening and e’ increase). This is consistent with systo-diastolic coupling, i.e. the contribution to early relaxation of elastic restitution (recoil) of systolic force (Burns et al., 2009).

Beside affording inotropy and lusitropy, SERCA2a stimulation may improve intracellular Ca^2+^ compartmentalization, with potential long-term effects on energetic efficiency and biology of cardiac myocytes (Zaza and Rocchetti, 2015).

PST3093 is remarkably less toxic than istaroxime which, in turn, has a lower proarrhythmic risk as compared to digoxin (Micheletti et al., 2002). We speculate that the low PST3093 toxicity, relative to istaroxime, may be due its failure to inhibit the Na^+^/K^+^ pump. Absence of interaction with 50 cardiac and non-cardiac targets commonly involved in drug toxicity provides at least a first level evidence of PST3093 suitability as a therapeutic agent.

### Limitations

Whereas, in the *in-vivo* experiments PST3093 effects generally achieved a maximum at 30 min of infusion, istaroxime infusion period was limited to 15 min. Our previous study (Torre et al., 2021) indicates that a 15 min infusion is sufficient for modulation of diastolic parameters by istaroxime. This time-point was selected in the present study to minimize metabolism to PST3093, thus allowing it to differentiate istaroxime’s own effect from that of its metabolite. Nonetheless, istaroxime effects reported here might differ from the steady-state ones, described in previous studies (Carubelli et al., 2020; Sabbah et al., 2007; Shah et al., 2009), to which PST3093 (the metabolite) might actually contribute.

Translation of the present *in-vivo* results to human therapy has to consider differences between clinical HF and the STZ rat model, which has specific hemodynamic features (Mihm et al., 2001). However, consistency of the istaroxime effect reported here with that described in HF patients (Carubelli et al., 2020) supports this translation.

### Therapeutic relevance and perspective

The results of this study identify PST3093 as a prototype “selective” (i.e. devoid of Na^+^/K^+^ pump inhibition) SERCA2a activator. This may entail significant differences from the already characterized pharmacodynamic profile of istaroxime.

In the case of istaroxime, lack of the proarrhythmic effect (expected from Na^+^/K^+^ ATPase inhibition (Rocchetti et al., 2005)) is likely due to SERCA2a stimulation. Indeed, the latter may reduce the occurrence of “Ca^2+^ waves” and the resulting “triggered activity” (Bai et al., 2013; Fernandez-Tenorio and Niggli, 2018; Zaza and Rocchetti, 2015). It is logical to predict that a pure SERCA2a activator may exert substantial antiarrhythmic effects, at least under the common conditions characterized by SR instability (e.g. HF). On the other hand, Na^+^/K^+^ pump inhibition may contribute to inotropy; thus, at least theoretically, PST3093 should increase systolic force less than istaroxime. The present results argue for a PST3093 effect on global cardiac function, including positive inotropy. Moreover, compared to istaroxime, PST3093 has a much longer half-life that, per se, may also prolong the beneficial hemodynamic effect of istaroxime infusion.

## Conclusions

HF treatment would strongly benefit from the availability of ino-lusitropic agents with a favorable profile. PST3093 is the main metabolite of istaroxime showing a longer half-life in human circulation compared to parent drug, activates SERCA2a, doesn’t inhibit Na^+^/K^+^ ATPase and improves systolic and diastolic performance in a model of diabetic cardiomyopathy. Overall, PST3093 acting as “selective” SERCA2a activator can be considered the prototype of a novel pharmacodynamic class for the ino-lusitropic approach of HF. After more than 50 years from the suggestion of the involvement of a reduced SERCA2a function as a cause of the depressed cardiac function and the increased arrhythmias in HF, we may have the possibility to prove this hypothesis and provide a “causal” and selective therapy for HF patients.

## Additional information

### Author Contributions

Conceptualization, investigation, formal analysis MA and MF; investigation and data curation PB, S-CH and ET; formal analysis AL and CR; resources and supervision G-JC, PF and GB; validation and funding acquisition FP; project administration, validation, supervision, funding acquisition, writing-original draft MR; project administration, validation, supervision, funding acquisition, writing – review and editing AZ.

### Additional file

Supplementary methods, Figures and Tables.

