## Supplementary material for "Istaroxime metabolite PST3093 selectively stimulates SERCA2a and reverses disease-induced changes in cardiac function"

by Arici M & Ferrandi M et al.

### Supplementary methods

*Animal models*

Male Sprague Dawley (SD) rats (150-175 gr) were used to generate STZ-induced diabetic cardiomyopathy model to test compounds *in vivo* and *in vitro*; female Dunkin-Hartley guinea pigs (175-200 g) were used for cardiac and skeletal muscle preparations (male 450-500 g) and myocytes isolation and finally, male Albino Swiss CD1 mice (30 g) were used for acute *in vivo* toxicity.

*Biochemical measurements*

Renal $Na^+/K^+$ ATPase purification and activity. Purification of renal $Na^+/K^+$ ATPase was performed according to the method of Jørgensen (Jørgensen, 1988). Frozen kidneys from 1-3 years-old male beagle dogs were obtained from the General Pharmacology Department of Sigma-tau, Pomezia, Italy. Kidneys were sliced and the outer medulla was dissected, pooled and suspended in a sucrose-histidine solution, containing 250 mM sucrose, 30 mM histidine and 5 mM EDTA, pH 7.2 and homogenized. The homogenate was centrifuged at 6,000 g for 15 min, the supernatant was decanted and centrifuged at 48,000 g for 30 min. The pellet was suspended in the sucrose-histidine buffer and incubated for 20 min with a sodium-dodecyl-sulphate (SDS) solution, dissolved in a gradient buffer, containing 25 mM imidazole and 1 mM EDTA, pH 7.5. The sample was layered on the top of a sucrose discontinuous gradient (10, 15 and 29.4%) and centrifuged at 60,000 g for 115 min. The final pellet was suspended in the gradient buffer. $Na^+/K^+$ ATPase activity was assayed by measuring $^{32}P$-ATP hydrolysis, as previously described (Ferrandi et al., 1996). Increasing concentrations of the standard ouabain, or tested compound, were incubated with 0.3 µg of purified dog kidney enzyme for 10 min at 37°C in 120 µl final volume of a medium, containing 140 mM NaCl, 3 mM $MgCl_2$, 50 mM Hepes-Tris, 3 mM ATP, pH 7.5. Then, 10 µl of incubation solution containing 10 mM KCl and 20 nCi of $^{32}P$-ATP (3-10 Ci/mmol, Perkin Elmer) were added, the reaction continued for 15 min at 37°C and was stopped by acidification with 20% ice-cold perchloric acid. $^{32}P$ was separated by centrifugation with activated Charcoal (Norit A, Serva) and the radioactivity was measured. Effects of increasing concentrations of the test compound were compared to ouabain, as positive standard, and to vehicle (control) at 37°C. The inhibitory activity was expressed as percent of activity in control.

SERCA ATPase activity assay. LV were dissected from rat and guinea pig of healthy and failing preparations and frozen until use. Tissues were homogenized, subjected to centrifugation to obtain SR-enriched microsomes and sarcomeric proteins were extracted, as previously described (Micheletti et al., 2007). In the case of rat hearts, cardiac homogenates were used in order to have sufficient material to replicate the experiments within a single animal. LV tissues from healthy and STZ rats were homogenized in a medium containing 300 mM sucrose, 50 mM K-phosphate, 10 mM NaF, 0.3 mM PMSF, 0.5 mM DTT (pH 7) and centrifuged at 35,000 g for 30 min. The final pellet was resuspended in the same buffer. In the case of guinea-pig hearts, LV tissues were homogenized in 4 volumes of 10 mM $NaHCO_3$, 1 mM PMSF, 10 µg/ml aprotinin and leupeptin (pH 7) and centrifuged at 12,000g for 15 minutes. Supernatants were filtered and centrifuged at 100,000 g for 30 min. Contractile proteins were extracted by suspending the pellets with 0.6 M KCl, 30 mM histidine, pH 7 and further centrifugation at 100,000 g for 30 min. Final pellets were reconstituted with 0.3 M sucrose, 30 mM histidine, pH 7, to obtain SR-enriched microsomes. SERCA2a ATPase activity was measured as the rate of $^{32}$P-ATP release at multiple $Ca^{2+}$ concentrations in the absence and presence of test compounds, as previously described (Micheletti et al., 2007). Increasing concentrations of each compound were pre-incubated with 2 µg of cardiac preparations for 5 min at 4° C in 80 µl of a solution containing 100 mM KCl, 5 mM $MgCl_2$, 1 µM A23187, 20 mM Tris, pH 7.5. Then, 20 µl of 5 mM Tris-ATP containing 50 nCi of $^{32}$P-ATP (3-10 Ci/mmol, Perkin Elmer) were added. The ATP hydrolysis was continued for 15 min at 37°C and the reaction was stopped by acidification with 100 µl of 20% ice-cold perchloric acid. $^{32}$P was separated by centrifugation with activated charcoal (Norit A, Serva) and the radioactivity was measured.

Reconstitution of SERCA1 with $PLN_{1-32}$ synthetic fragment. Adult healthy guinea-pigs were used to prepare SERCA1-enriched SR microsomes from fast-twitch hind leg muscles. Microsomes were prepared as described for SERCA2a preparations. For reconstitution experiments, SERCA1 (PLN-free) was pre-incubated with synthetic $PLN_{1-32}$ fragment (canine sequence, Biomatik Corporation, Canada) in 20 mM imidazole, pH 7, at 1:300 SERCA1:PLN ratio for 30 min at room temperature. After pre-incubation, SERCA1 (PLN-free) alone, or reconstituted with $PLN_{1-32}$ fragment, was utilized for SERCA activity measurement by using $^{32}$P-ATP hydrolysis method at different $Ca^{2+}$ concentrations in the absence and presence of increasing concentrations of tested compounds, as described for SERCA2a ATPase activity.

*PST3093 interaction with targets other than SERCA*

Analysis of PST3093 interaction with a panel of 50 ligands was carried out by Eurofins (Taiwan) on crude membrane preparations according to Eurofins described procedures. The assay is partly based on radioligand displacement (e.g. for receptors) and partly on spectrophotometric detection of change in function (e.g. for enzymes). Results were compared to appropriate reference standards; a >50% change in affinity or activity was considered as a positive hit (interaction present).

*Functional measurements in isolated myocytes*

Rat and guinea-pig LV ventricular myocytes were isolated by using a retrograde coronary perfusion method previously published (Rocchetti et al., 2003) with minor modifications. Rod-shaped, $Ca^{2+}$-tolerant myocytes were used within 12 h from dissociation. LV myocytes were clamped in the whole-cell configuration (Axopatch 200A, Axon Instruments Inc., Union City, CA). During measurements, myocytes were superfused at 2 ml/min with Tyrode's solution containing 154 mM NaCl, 4 mM KCl, 2 mM $CaCl_2$, 1 mM $MgCl_2$, 5 mM HEPES/NaOH, and 5.5 mM D-glucose, adjusted to pH 7.35. A thermostated manifold, allowing for fast (electronically timed) solution switch, was used for cell superfusion. All measurements were performed at 35 °C. The pipette solution contained 110 mM

$K^+$ -aspartate, 23 mM KCl, 0.2 mM $CaCl_2$ ($10^{-7}$ M calculated free-$Ca^{2+}$ concentration), 3 mM $MgCl_2$, 5 mM HEPES-KOH, 0.5 mM EGTA-KOH, 0.4 mM GTP-$Na^+$ salt, 5 mM ATP-$Na^+$ salt, and 5 mM creatine phosphate $Na^+$ salt, pH 7.3. Membrane capacitance and series resistance were measured in every cell but left uncompensated. Current signals were filtered at 2 KHz and digitized at 5 KHz (Axon Digidata 1200). Trace acquisition and analysis was controlled by dedicated software (Axon pClamp 8.0).

$Na^+/K^+$ ATPase current ($I_{NaK}$) measurements. $I_{NaK}$ was recorded in isolated rat LV myocytes (Alemanni et al., 2011; Rocchetti et al., 2003) as the holding current recorded at -40 mV in the presence of $Ni^{2+}$ (5 mM), nifedipine (5 µM), $Ba^{2+}$ (1 mM) and 4-aminopyridine (2 mM) to minimize contamination by changes in $Na^+/Ca^{2+}$ exchanger (NCX), $Ca^{2+}$ and $K^+$ currents, respectively. Tetraethylammonium-Cl (20 mM) and EGTA (5 mM) were added to the pipette solution and intracellular $K^+$ was replaced by $Cs^+$. To optimize the recording conditions, $I_{NaK}$ was enhanced by increasing intracellular $Na^+$ (10 mM) and extracellular $K^+$ (5.4 mM). All drugs were dissolved in dimethyl sulfoxide (DMSO). Control and test solutions contained maximum 1:100 DMSO.

Intracellular $Ca^{2+}$ dynamics. LV myocytes were incubated in Tyrode's solution for 30 min with the membrane-permeant form of the dye, Fluo4-AM (10 µM), and then washed for 15 min to allow dye de-esterification. Fluo4 emission was collected through a 535 nm band pass filter, converted to voltage, low-pass filtered (100 Hz) and digitized at 2 kHz after further low-pass digital filtering (FFT, 50 Hz). After subtraction of background luminescence, a reference fluorescence ($F_0$) value was used for signal normalization ($F/F_0$). Cytosolic $Ca^{2+}$ activity was dynamically measured in field stimulated (2 Hz) and patch-clamped rat LV myocytes. In the first case, fluorescence in diastole was used as $F_0$ for signal normalization ($F/F_0$).

In patch-clamped myocytes membrane current, whose time-dependent component mainly reflected the sarcolemmal $Ca^{2+}$ current ($I_{CaL}$), was simultaneously recorded. Drug effects on SR $Ca^{2+}$ uptake rate were evaluated with a "SR loading protocol" specifically devised to rule out the contribution of NCX and to assess the SR $Ca^{2+}$ uptake rate at multiple levels of SR $Ca^{2+}$ loading (Rocchetti et al., 2005) (protocol in Figure S1). The protocol consisted in emptying the SR by a brief caffeine (10 mM) pulse and then progressively refilling it by 7-10 voltage steps (-35 to 0 mV) activating $Ca^{2+}$ influx through $I_{CaL}$. NCX was blocked by omission of $Na^+$ from intracellular and extracellular (replaced by equimolar $Li^+$ and 1 mM EGTA) solutions. The procedure is in agreement with published methods, with minor modifications (Alemanni et al., 2011; Rocchetti et al., 2005; Torre et al., 2021). Multiple parameters, suitable to quantify SR $Ca^{2+}$ uptake, can be extracted from $Ca^{2+}$ and $I_{CaL}$ response to the protocol: the time constant ($\tau$) of cytosolic $Ca^{2+}$ decay within each V-step largely reflects net $Ca^{2+}$ flux across the SR membrane (the faster SR $Ca^{2+}$ uptake, the smaller $\tau$ decay). Because of the steep dependency of CaT amplitude on SR $Ca^{2+}$ content, the rate at which CaT amplitude increases across the subsequent pulses of the protocol reflects the rate at which the SR refills. To rule out the potential contribution of changes in $I_{CaL}$, in each loading step, CaT amplitude was normalized to $Ca^{2+}$ influx (estimated from $I_{CaL}$ integral up to CaT peak) to obtain excitation-release (ER) gain. As expected from its strong dependency on SR $Ca^{2+}$ content, this parameter progressively increases during the loading protocol. Diastolic $Ca^{2+}$ of the first step was used as $F_0$ for signal normalization ($F/F_0$). Specificity of the "loading protocol" parameters in detecting SERCA2a activation is supported by the observation that they did not detect any effect of digoxin, an inotropic agent blocking the $Na^+/K^+$ pump and devoid of SERCA2a stimulating effect (Alemanni et al., 2011; Rocchetti et al., 2005).

**Supplementary Figures and Tables:**

- Figure S1: Protocol to evaluate intracellular $Ca^{2+}$ dynamics in patch-clamped cells under $Na^+$ free condition;
- Figure S2: Analytical characterization of PST3093;
- Figure S3: Effects of 1 µM PST3093 on electrical activity in guinea-pig myocytes;
- Table S1: Effect of PST3093 and istaroxime on SERCA2a kinetic parameters in cardiac preparations from healthy guinea-pigs;
- Table S2: Effect of PST3093 (10 µM) on the panel of molecular targets; Table S3: Characterization of the STZ diabetic rat model; Table S4: Echocardiographic parameters in healthy and STZ diabetic rats.

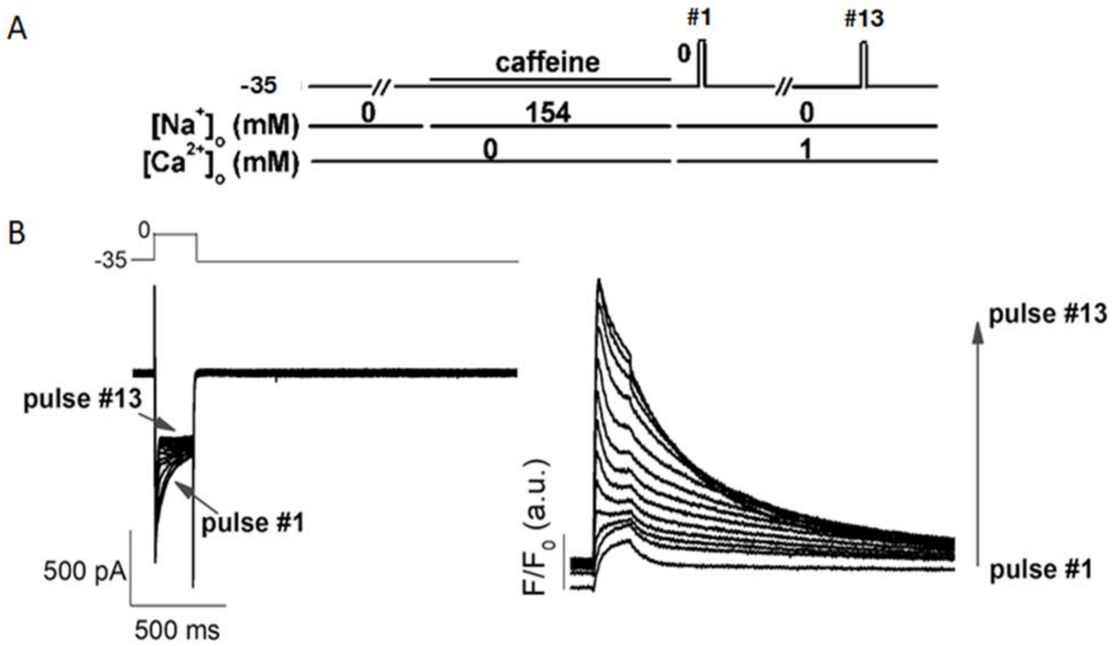


**Figure S1. Protocol to evaluate intracellular $Ca^{2+}$ dynamics in patch-clamped cells under $Na^{+}$ free condition**. **A)** Protocol outline. **B)** Transmembrane current (left) and $Ca^{2+}$ transients (right) recordings during SR reloading after caffeine-induced SR depletion in patch-clamped cells. See Methods for details.

**A** $^{1}$H-NMR

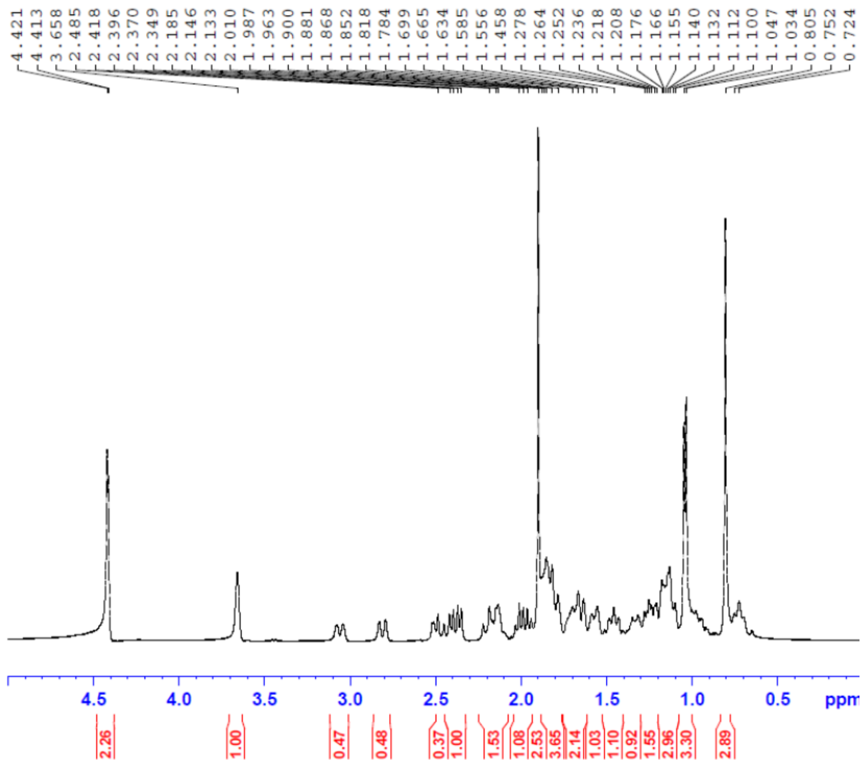


**B** $^{13}$C-NMR

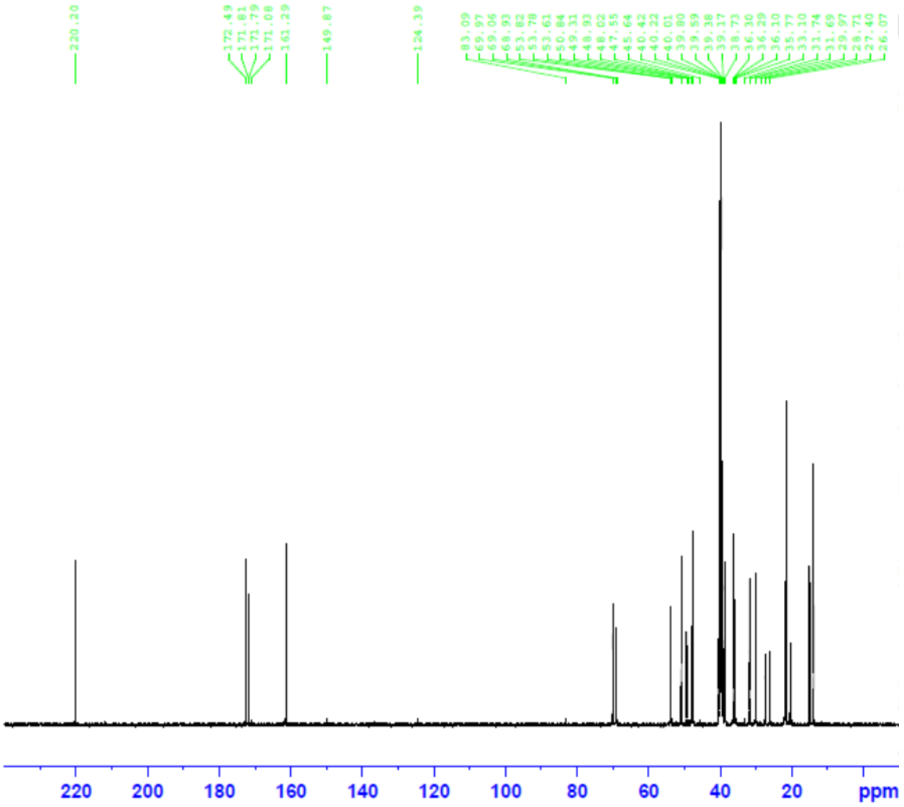

**C**

Sample Information

| | | | |
|---|---|---|---|
| Injection Date: | 6/17/2015 | Acq.Operator: | spc |
| Sample Name: | 0505-021-06-PST3093 | Acq. Method: | D:\DATA\SPC3374\ -> |
| Location: | Vial 21 | Injection Time: | 7:02:00 PM |
| Injection#: | 1 | Injection Volume: | 10.0 uL |
| Seq. Line: | 7 | Sample Amount: | 0 mg/ml |
| Run Time: | 30min | | |

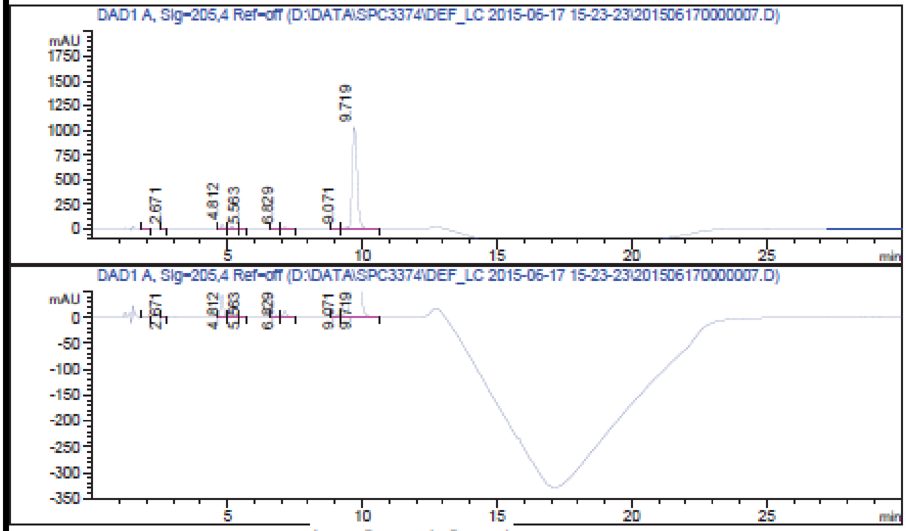


Area Percent Report

Signal: DAD1 A, Sig=205,4 Ref=off

| # | Compound Name | Meas. R | Peak T | Width | Area | Height | Area % |
|---|---|---|---|---|---|---|---|
| 1 | | 2.081 | MM | 0.116 | 29.910 | 4.293 | 0.211 |
| 2 | | 2.671 | BV | 0.085 | 5.830 | 1.052 | 0.041 |
| 3 | | 4.812 | BV | 0.121 | 338.417 | 43.471 | 2.384 |
| 4 | | 5.164 | VV | 0.118 | 174.666 | 22.747 | 1.231 |
| 5 | | 5.563 | VB | 0.122 | 9.090 | 1.176 | 0.064 |
| 6 | | 6.829 | BV | 0.142 | 31.993 | 3.441 | 0.225 |
| 7 | | 7.133 | VB | 0.151 | 125.430 | 12.685 | 0.884 |
| 8 | | 9.071 | BV | 0.165 | 52.584 | 4.919 | 0.371 |
| 9 | | 9.719 | VB | 0.201 | 13424.721 | 1.034e3 | 94.589 |

*** End of Report ***

Operator: Review:

**Figure S2. Analytical characterization of PST3093. A)** $^{1}$H-NMR in DMSO (400 MHz, Bruker), **B)** $^{13}$C-NMR in DMSO (100 MHz, Bruker), **C)** HPLC profile (about 95% purity).

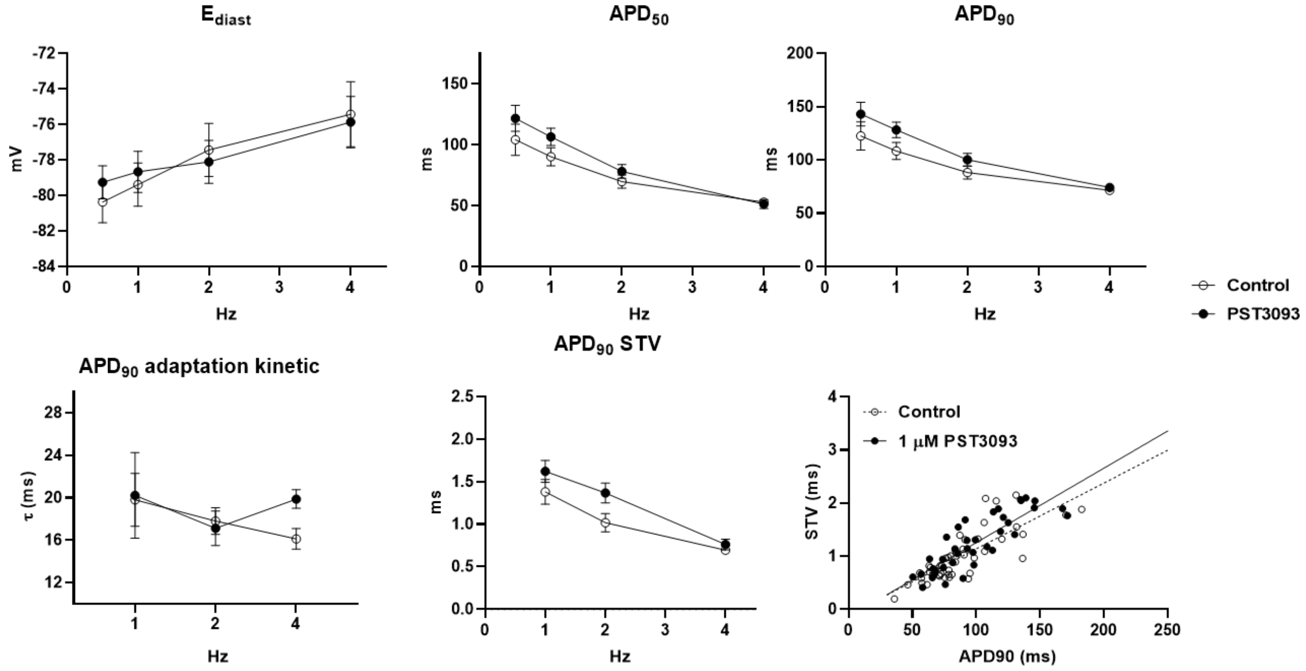


**Figure S3. Effects of 1 µM PST3093 on electrical activity in guinea-pig myocytes.** Rate dependency of AP parameters ($E_{diast}$, $APD_{50}$, $APD_{90}$), $APD_{90}$ adaptation kinetics and $APD_{90}$ short term variability (STV) with or w/o 1 µM PST3093; N=3 (n=16 w/o PST3093, n=18 with PST3093). Bottom right: linear correlation between STV of $APD_{90}$ and $APD_{90}$ values with or w/o PST3093; data from 1, 2, 4 Hz were pooled (w/o PST3093 slope = 0.012, with PST3093 slope = 0.014, NS).

**Table S1. Effect of PST3093 and istaroxime on SERCA2a kinetic parameters in cardiac preparations from healthy guinea-pigs.** Data are mean ± SEM; N indicates the number of experiments. *$p<0.05$ vs control (RM one-way ANOVA with post-hoc Tukey's multiple comparisons test or paired *t*-test).

| Guinea-pigs | | | | | | |
|---|---|---|---|---|---|---|
| **[nM]** | **N** | **control** | **PST3093** | **N** | **control** | **istaroxime** |
| **$V_{max}$ (µmol/min/mg)** | | | | | | |
| **1** | 9 | 2,28±0,17 | 2,16±0,18 | 9 | 2,51±0,08 | 2,50±0,04 |
| **10** | | | 2,15±0,15 | | | 2,47±0,06 |
| **100** | 11 | 2,38±0,16 | 2,34±0,16 | | | 2,49±0,07 |
| **$K_dCa$ (nM)** | | | | | | |
| **1** | 9 | 567±12 | 504±23* | 9 | 584±22 | 524±19 |
| **10** | | | 501±17* | | | 485±16* |
| **100** | 11 | 557±12 | 445±30* | | | 478±19* |

**Table S2. Effect of PST3093 (10 µM) on the panel of molecular targets** (Eurofins, Taiwan).

| | Cat # | Assay name | Batch | Species | PST3093 effect (%) |
|---|---|---|---|---|---|
| 1 | 107480 | ATPase, $Ca^{2+}$, skeletal muscle | 438642 | pig | -18 |
| 2 | 118040 | CYP450, 19 | 438644 | human | 12 |
| 3 | 124010 | HMG-CoA Reductase | 438610 | human | -2 |
| 4 | 140010 | Monoamine Oxidase MAO-A | 438645 | human | 2 |
| 5 | 140120 | Monoamine Oxidase MAO-B | 438647 | human | -3 |
| 6 | 143000 | Nitric Oxide Synthase, Endothelial (eNOS) | 438568 | bovine | 2 |
| 7 | 107300 | Peptidase, Angiotensin Converting Enzyme | 438641 | rabbit | 3 |
| 8 | 164610 | Peptidase, Renin | 438648 | human | 6 |
| 9 | 152000 | Phosphodiesterase PDE3 | 438611 | human | -3 |
| 10 | 171601 | Protein Tyrosine Kinase, ABL1 | 438612 | human | 3 |
| 11 | 176810 | Protein Tyrosine Kinase, Src | 438613 | human | -1 |
| 12 | 200510 | Adenosine A1 | 438614 | human | -2 |
| 13 | 200610 | Adenosine A2A | 438614 | human | -6 |
| 14 | 203100 | Adrenergic α1A | 438615 | rat | 2 |
| 15 | 203200 | Adrenergic α1B | 438615 | rat | 0 |
| 16 | 203630 | Adrenergic α2A | 438616 | human | -5 |
| 17 | 204010 | Adrenergic β1 | 438652 | human | -4 |
| 18 | 204110 | Adrenergic β2 | 438571 | human | 7 |
| 19 | 204600 | Aldosterone | 438617 | rat | 5 |
| 20 | 206000 | Androgen (Testosterone) | 438618 | human | 3 |
| 21 | 210030 | Angiotensin AT1 | 438653 | human | -1 |
| 22 | 210120 | Angiotensin AT2 | 438653 | human | 7 |
| 23 | 214600 | Calcium Channel L-type, Dihydropyridine | 438620 | rat | -8 |
| 24 | 219500 | Dopamine D1 | 438660 | human | 3 |
| 25 | 219700 | Dopamine D2s | 439024 | human | 26 |
| 26 | 219800 | Dopamine D3 | 438660 | human | 1 |
| 27 | 226010 | Estrogen ERα | 438622 | human | -6 |
| 28 | 226050 | Estrogen ERβ | 438622 | human | 7 |

| | | | | | |
|---|---|---|---|---|---|
| 29 | 226600 | $GABA_A$, Flunitrazepam, Central | 438624 | rat | 4 |
| 30 | 226500 | $GABA_A$, Muscimol, Central | 438623 | rat | 9 |
| 31 | 232030 | Glucocorticoid | 438626 | human | 8 |
| 32 | 233000 | Glutamate, NMDA, Phencyclidine | 438627 | rat | -8 |
| 33 | 239610 | Histamine H1 | 438628 | human | -3 |
| 34 | 241000 | Imidazoline I2, Central | 438629 | rat | 5 |
| 35 | 243000 | Insulin | 438654 | rat | 6 |
| 36 | 252710 | Muscarinic M2 | 438621 | human | -7 |
| 37 | 252810 | Muscarinic M3 | 438661 | human | 4 |
| 38 | 253010 | Muscarinic M5 | 438661 | human | 5 |
| 39 | 258730 | Nicotinic Acetylcholine α3β4 | 438656 | human | 1 |
| 40 | 260410 | Opiate μ (OP3, MOP) | 438616 | human | -5 |
| 41 | 264500 | Phorbol Ester | 438624 | mouse | -7 |
| 42 | 265600 | Potassium Channel ($K_{ATP}$) | 438632 | hamster | -8 |
| 43 | 265900 | Potassium Channel hERG | 438633 | human | 10 |
| 44 | 299005 | Progesterone PR-B | 438638 | human | -15 |
| 45 | 270300 | Ryanodine RyR3 | 438634 | rat | 2 |
| 46 | 271010 | Serotonin (5-Hydroxytryptamine) 5-HT1, non-selective | 438668 | rat | 0 |
| 47 | 299007 | Sigma σ2 | 438662 | human | 12 |
| 48 | 278110 | Sigma σ1 | 438636 | human | 7 |
| 49 | 279510 | Sodium Channel, Site 2 | 438637 | rat | 2 |
| 50 | 204410 | Transporter, Norepinephrine (NET) | 438597 | human | 4 |

**Table S3. Characterization of the STZ diabetic rat model**. Parameters were measured after 1 or 8 weeks (wk) from STZ injection. Data are mean ± SEM. N represents the number of rats for each group. *p<0.05 vs healthy rats (unpaired *t*-test).

| | **Healthy** | | **STZ** | |
|---|---|---|---|---|
| **Parameters** | **1 wk after vehicle** | **8 wk after vehicle** | **1 wk after STZ** | **8 wk after STZ** |
| BW (g) | 201±4.7 | 449±11.8 | 193±4.2 | 305±12.9* |
| Glycaemia (mg/dL) | 130±6.6 | nd | 398±16* | nd |
| LV mass ($mm^3$) | nd | 985±34 | nd | 781±40* |
| LV mass index ($mm^3$/g) | nd | 2,23±0,1 | nd | 2,66±0,2 |
| N | 18 | 18 | 20 | 20 |

**Table S4. Echocardiographic parameters in healthy and STZ diabetic rats**. Data are mean ± SEM. N represents the number of rats for each group. *p<0.05 vs healthy rats (unpaired *t*-test).

| **Echo parameters** | | **Healthy** | **STZ** |
|---|---|---|---|
| **Morphometric parameters** | IVSTd (mm) | 2,22±0,07 | 1,73±0,1 * |
| | PWTd (mm) | 1,87±0,09 | 1,54±0,05 * |
| | LVEDD (mm) | 6,6±0,14 | 6,95±0,13 |
| | IVSTs (mm) | 2,46±0,07 | 2,26±0,1 |
| | PWTs (mm) | 2,96±0,09 | 2,47±0,11 * |
| | LVESD (mm) | 3,09±0,09 | 3,54±0,1 * |
| **Systolic parameters** | FS (%) | 53,08±1,17 | 48,73±1,37 * |
| | s' (mm/s) | 29,59±1,15 | 24,8±0,75 * |
| | EF (%) | 88 ± 0,8 | 85 ±1,08 * |
| **Diastolic parameters** | E (mm/s) | 0,93±0,01 | 0,84±0,02 * |
| | A (mm/s) | 0,65±0,04 | 0,63±0,03 |
| | E/A | 1,50±0,08 | 1,38±0,07 |
| | DT (ms) | 54,61±2,1 | 56,8±2,16 |
| | DT/E | 59,23±2,4 | 68,37±3,0 * |
| | E/DT | 17,41±0,77 | 15,18±0,67 * |
| | e' (mm/s) | 23,46±0,77 | 21,34±0,58 * |
| | a' (mm/s) | 24,54±1,38 | 24,3±1,13 |
| | e'/a' | 1,01±0,06 | 0,91±0,04 |
| | E/e' | 40,1±1,33 | 39,64±1,0 |
| **Overall cardiac function** | HR (bpm) | 306±10 | 248±7,4 * |
| | SV (ml) | 0,59±0,03 | 0,65±0,04 |
| | CO (ml/min) | 178,9±10,2 | 161,1±10,9 |
| | N | 18 | 20 |
